# Blood-meal analysis indicates preferential feeding on humans and non-commensal rodents by *Culex* mosquitoes in Tanzania

**DOI:** 10.64898/2026.11.10.765634

**Authors:** Wim De Kesel, Robin Fierens, Baraka E. Mwamundela, Abdul S. Katakweba, Joachim Mariën, Erik Verheyen, Kevin K. Ariën, Sophie Gryseels

## Abstract

Arthropod-borne viruses are important emerging and re-emerging pathogens worldwide, yet information on mosquito feeding ecology and associated transmission networks remains limited in many African regions. This study investigated mosquito host-feeding patterns and arbovirus occurrence along an urban-rural gradient in Morogoro, Tanzania. Mosquitoes were collected at four locations representing different levels of human influence during wet and dry seasons. Vertebrate blood-meal sources were identified using a Nanopore-based metabarcoding approach, while viral screening was performed using RT-qPCR assays targeting *Orthoflavivirus* and *Alphavirus* genera.

A total of 8,917 mosquitoes were collected, of which more than 90% belonged to the *Culex* genus. Blood-meal analyses of 310 blood-fed and gravid mosquitoes identified 35 molecular operational taxonomic units, which we grouped into 11 host categories. Humans were the most frequently utilized hosts (68%), followed by non-commensal rodents (22%) and crows (18%). *Culex* mosquitoes showed broad host use and frequently fed on multiple taxonomic units, with an average of 1.36 taxonomic units detected per individual. Co-feeding network analyses revealed extensive host sharing between humans and wildlife, with non-commensal rodents showing significantly higher co-feeding probabilities with humans than any other host group. No human-pathogenic arboviruses were detected. Viral detections were limited to insect-specific viruses, an endogenous viral element in chromosome 2 of *Aedes aegypti*, and two detections of bird-infecting Bagaza virus, indicating ongoing viral circulation within the mosquito community.

Our findings demonstrate broad and flexible host use by *Culex* mosquitoes and highlight non-commensal rodents as an underappreciated component of mosquito feeding with potential relevance for arbovirus transmission and spillover between wildlife and humans.

## Introduction

Mosquitoes are the most common vectors of arboviruses, with several genera – including *Aedes*, *Culex*, and *Anopheles* - capable of transmitting medically relevant viruses [1,2]. Through their hematophagous behaviour mosquitoes acquire and transmit viruses while feeding on vertebrate hosts, with the range of hosts varying according to species-specific feeding preferences [1,3]. In addition, many mosquito species exhibit strong associations with specific environments, with some being highly adapted to human-modified landscapes [4,5]. Life-history traits, such as short generation times and high reproductive capacity, combined with extrinsic factors, such as temperature and humidity, facilitate rapid population growth and sustained virus transmission [6–9].

Arbovirus transmission dynamics are shaped not only by the presence of competent vectors but also by interactions between mosquitoes and vertebrate hosts. Mosquito host selection is influenced by a combination of host abundance, accessibility, habitat overlap, and species-specific feeding preferences, resulting in considerable variation in host use among mosquito species [10–15]. Because host-feeding patterns determine which vertebrate species are exposed to mosquito bites and how frequently mosquitoes feed across different host taxa, they directly influence pathogen circulation, the involvement of vertebrate hosts in transmission cycles, and the risk of spillover to humans [16–19]. Vertebrate hosts further differ in their susceptibility to infection and their capacity to amplify and transmit arboviruses, although the role of many host species remains poorly understood [16–18]. Consequently, identifying mosquito blood-meal sources provides valuable insight into the ecological networks that sustain arbovirus transmission and can help identify potential reservoir hosts, bridge hosts, and pathways of pathogen emergence [20,21].

Studies across eastern Africa have demonstrated that mosquitoes feed on a remarkably diverse range of vertebrate hosts, including humans, livestock, birds, reptiles, amphibians, and wildlife, although the relative importance of these hosts varies among mosquito species and ecological settings. Livestock, particularly cattle, goats, and sheep, are among the most frequently identified blood-meal sources and often constitute a substantial proportion of mosquito blood meals in rural and pastoral landscapes [22–26]. Human blood meals are also common, especially among anthropophilic species such as *Aedes* spp. and *Anopheles* spp. [22,25,26]. Beyond domestic hosts, mosquitoes have been reported feeding on a wide diversity of vertebrates, including dogs, pigs, rodents, bats, non-human primates, amphibians, and numerous wild bird species [23,24]. Studies conducted at wildlife-livestock interfaces have further demonstrated that individual mosquito species frequently feed across wildlife, livestock, and human host groups, indicating substantial ecological connectivity among vertebrate communities [23]. Recent molecular approaches have additionally revealed that mixed blood meals are common, with individual mosquitoes often feeding on multiple host species within a single gonotrophic cycle [24–26]. Such opportunistic feeding patterns may facilitate pathogen maintenance in wildlife and livestock populations while simultaneously creating opportunities for transmission to humans.

Despite growing evidence that eastern African mosquitoes utilize a broad diversity of vertebrate hosts, information remains geographically incomplete and is particularly scarce for urban and peri-urban ecosystems, limiting our understanding of the host communities involved in arbovirus transmission. Improved characterization of mosquito feeding patterns, complemented by arbovirus surveillance, can help identify vertebrate species that are frequently connected through mosquito feeding interactions and may contribute to local transmission cycles.

This study aimed to characterize mosquito blood-host diversity and identify arboviruses carried by mosquitoes along an urban-rural gradient in Tanzania, a region where several arboviruses, such as Chikungunya virus, Rift Valley fever virus, West Nile virus, and dengue virus, have caused outbreaks in recent decades [27–31]. We addressed three research question: Which vertebrate hosts are most frequently utilized by mosquitoes? To what extent do mosquitoes feed on multiple host species, thereby creating potential pathways for pathogen transmission? Are there important mosquito hosts that have been overlooked in arbovirus transmission studies? We hypothesize that mosquitoes feed preferentially on certain vertebrate species, and as such that the risk of arbovirus transmission from non-human hosts to humans is skewed towards originating from these preferred vertebrate species.

## Material & Methods

### - Study area and sampling design

Sampling was conducted in Tanzania in the city of Morogoro and its surrounding areas (Figure 1). Four sites were selected representing an urban-to-rural gradient, one urban site in the city centre, a suburban site in the transition zone of urban to agricultural area and two rural sites in the transition zone of agricultural to forest area. Sampling was performed during the post-rainy season (May) in 2023 and the dry season (August) in 2025. Mosquitoes were trapped using BG-Pro traps in EVS-style configuration without LED lights with the BG-CO₂ generator (Biogents, Germany). At each site simultaneously two traps were deployed 10 m apart, 1 m above the ground and active for 72 hours. Traps were emptied daily in the morning (9:00-10:00 AM) and 24h CO2 bait was replenished in the evening (5:00-6:00 PM).

**Figure 1:**
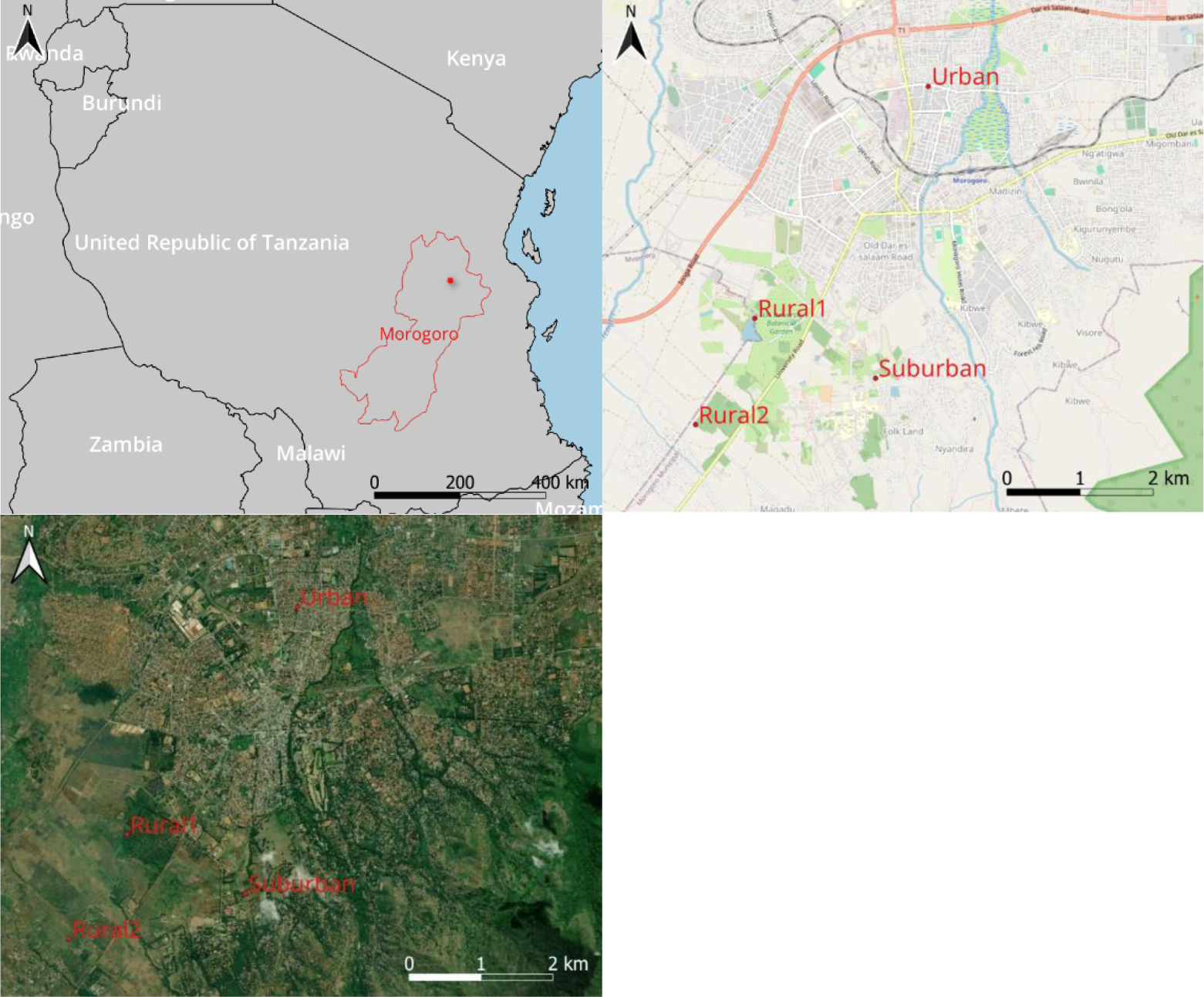
Maps of the study area. Top left: United Republic of Tanzania with a focus in red of the Morogoro region with the regional capital city, Morogoro, indicated with a red dot. Top right: detailed OpenStreetMap map of Morogoro city with the four sampling locations. Bottom left: detailed ESRI satellite map of Morogoro city with the four sampling locations. Maps create in QGIS 3.28.3, layers originate from QGIS, https://www.naturalearthdata.com/ and https://d-maps.com/.

### - Sample processing

Mosquitoes were euthanized by freezing at −20 °C for 15 min and morphologically identified to genus level using pictorial keys [32–34]. Males were pooled (20 individuals per pool) by genus and site and stored dry at −20 °C. Females were stored individually in 200 µL DNA/RNA Shield at −20 °C, and their abdominal status (blood-fed, gravid, or empty) was recorded. Nucleic acids were extracted using the Quick-DNA/RNA Pathogen Miniprep Kit (Zymo Research) following manual homogenization, with 1% proteinase K and omission of β-mercaptoethanol. All blood-fed and gravid females were extracted individually, together with a stratified subset of 30 non-blood-fed females and 30 males. Remaining non-blood-fed females were extracted in pools of approximately 16 individuals grouped by genus and site. Eluates were stored at −80 °C.

### - Mosquito and host identification by Nanopore metabarcoding

Only DNA/RNA extracts of individually extracted specimens were analysed for mosquito and host identity. Mosquito identity was determined by a 659bp COX1 fragment using primers LCO1490 and HCO2198 [35]. The 15µL reaction mix contained 1.5µL of DNA, 1X Green GoTaq Flexi Buffer (Promega), 1.5mM MgCl2, 0.2mM dNTP, 0.4µM of each primer, 0.6 units of G2 Flexi DNA Polymerase and nuclease free water. Cycling conditions consisted of 94°C for 3min, five cycles consisting of 94°C, 45°C and 72°C each for 30s, 35 cycles consisting of 94°C, 51°C, 72°C each for 30s, with a final elongation at 72°C for 5min. Host identity was based on a 108bp 12S rRNA gene fragment with primers 12S-V5f and 12S-V5r [36,37]. The reaction mixture contains the same ratios as the mosquito identity with the exception of 2.5mM MgCl2, 0.1µM of each primer and one unit of DNA Polymerase. Cycling conditions consisted of 95°C for 5min, five cycles consisting of 95°C, 45°C and 72°C each for 30s, 35 cycles consisting of 95°C, 53°C, 72°C each for 30s, with a final elongation at 72°C for 5min.

COX1 amplicons were quantified using a Qubit fluorometer (Thermo Fisher Scientific) and 100ng was pooled with the corresponding 12S amplicons. Pooled products were purified and eluted in 30µL using the NucleoSpin Gel and PCR Clean-up Kit (Macherey Nagel). Nanopore library was prepared using the Native Barcoding Kit 96 V14 (Oxford Nanopore Technologies), used protocol can be found in supplementary information (Nanopore_SamplePrep). Libraries were sequenced on MinION R10.4.1 flowcells for 24 hours. In brief, acquired reads were basecalled using the super accuracy model and filtered on quality (QC15) and length (>70bp). Consensus creations were generated using NGSpeciesID and taxonomical assignment was based on a curated mitochondrial database through BLASTN using the lowest common ancestor based on the bit score. Host and vector consensus sequences were corrected for background contamination by subtracting supporting reads observed in the corresponding extraction negative controls (or a default of 10 reads when absent). To account for barcode hopping and library cross-contamination, taxon-specific detection thresholds were calculated separately for hosts and vectors from PCR negative controls (maximum read count + 1 SD), with a minimum threshold of 100 supporting reads. Detailed information on the bioinformatics pipeline and consensus formation and filtering can be found in supplementary information: Bioinformatics.

### - Host analysis statistics

To retain only signals likely originating from the most recent blood meals, host sequences were filtered based on their proportional read abundance within each mosquito. The proportion threshold (13%) was empirically determined using 30 blood-fed, 30 non-blood-fed, and 30 male mosquitoes and calculated as the ratio between background host reads in controls and total host reads in blood-fed mosquitoes. Only host identifications exceeding this threshold were retained for downstream analyses. Host identifications were grouped into broader host categories based on the detected hosts to limit the variation in taxonomic level identifications.

The host identification data was converted into a binary matrix indicating the presence or absence of each host category within each mosquito specimen [38]. The McNemar test was done with holm adjusted p value to identify significant pairwise differences between host proportions [39]. Host–host association undirected weighted networks, using the *igraph* package, were based on shared mosquitoes between host groups using a Simple-Ratio Index (SRI) [38,40].

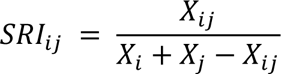

Where *Xij* is the number of mosquitoes shared between host *i* and *j*, and *Xi* and *Xj* are the total number of mosquitoes having fed on those hosts.

For blood meals containing human DNA, a generalized linear mixed-effects model (GLMM) with a binomial error distribution was used to test whether the probability of co-occurrence with humans differed among host categories. Host category was included as a fixed effect, and mosquito identifier was included as a random effect to account for repeated observations within blood meals. Season and site were not included because most samples originated from urban locations sampled during the wet season. Significance was assessed using likelihood ratio tests and estimated marginal means were calculated for significant variables, with pairwise comparisons. All data preparation, analysis and statistical procedures were conducted using R Statistical Software (R version 4.6.0) [41].

### - Arbovirus screening

All DNA/RNA extracts of individual mosquitoes and extracted pools were tested for *Orthoflavivirus* and *Alphavirus* using two one-step RT-qPCRs. *Orthoflavivirus* screening was targeted at a 230bp fragment of the NS5 gene using primers PF1S and PF2R-bis [42]. *Alphavirus* screening was targeted at a 179bp fragment of nsP4 gene using Alp-F and Alp-R [43]. Both 20µL qPCR reaction mixtures contained 5µL RNA, 10µL 2x SYBR-Green Master-Mix (QuantiTect SYBR-Green RT-PCR, Qiagen), 0.2µL RT-mix, 1µM of the respective forward and reverse primer and nuclease free water. Cycling conditions using a Light Cycler 480 (Roche) were: reverse transcription at 50°C for 30 minutes and activation at 95°C for 15 minutes; followed by 30 (*Orthoflavivirus*) or 40 (*Alphavirus*) cycles of 94°C for 15s, 50°C (*Orthoflavivirus*) or 45°C (*Alphavirus*) for 30s, and 72°C for 30s, with SYBR Green fluorescence acquisition. Melting curve analysis was then performed from 60°C to 95°C at 0.06°C/s with continuous fluorescence monitoring.

Blood-fed and gravid mosquito specimens with more than 50% domestic mammal or bird blood were screened for respectively Rift Valley fever virus (RVFV) or West Nile virus (WNV), by one-step RT-qPCR. Rift Valley fever virus screening was targeted at a 95bp G2 gene fragment using RVFV-fw and RVFV-rv [44]. West Nile virus detection was targeted at a 144bp fragment spanning the 5’ untranslated region and the capsid gene using ProC-F1 and ProC-R [45]. The reaction mixture ratios are the same as with *Orthoflavivirus* qPCR except for using 0.5µM of the primers. Cycling conditions were: reverse transcription at 50°C for 30 minutes and activation at 95°C for 15 minutes; 45 cycles at 94°C for 15s (WNV) or 5s (RVFV), 57°C for 15s, and 72°C for 15s, with SYBR Green fluorescence acquisition afterwards a melting curve analysis was performed.

Virus positivity was assessed based on the amplification and melting curve analyses. Potentially positive samples were Sanger sequencing in both directions at the Neuromics Support Facility of the VIB, Antwerp, Belgium. Acquired sequences were assembled using Geneious Prime’s de novo assembler with default settings (Geneious version 2026.1). Assembly consensus sequences containing <5% ambiguous bases were aligned with respective reference sequences and analysed using NCBI BLAST.

## Results

### - Captured mosquitoes

A total of 8,917 mosquitoes were collected across all sites (5,898 wet; 3,019 dry season). Most belonged to *Culex* (93.4%, 8,325), followed by *Aedes* (390), *Mansonia* (141), *Anopheles* (7), *Toxorhynchites* (6), and 46 unidentified specimens (Figure 2). Of all specimens, 4,949 were female (with 363 blood-fed or gravid), 3,907 male, and 61 of undetermined sex. Detailed counts are provided in the supplementary information (Mosquito_analysis).

**Figure 2:**
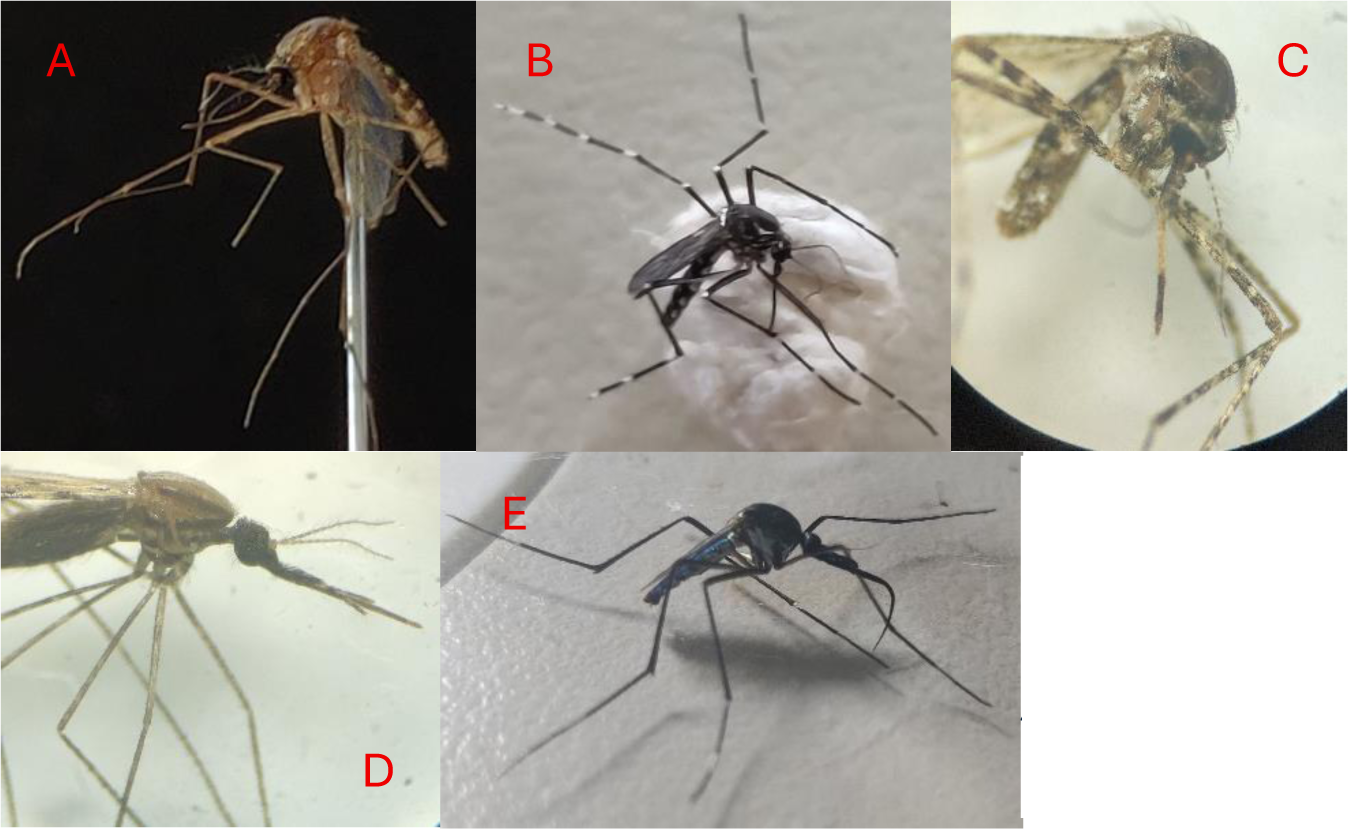
Representative pictures of captured mosquitoes. A: Female Culex pipiens pipiens, B: female Aedes aegypti, C: female Mansonia sp., D: female Anopheles sp. And E: female Toxorhynchites sp..

### - Mosquito and host identification

Blood meal results were acquired for 310 (85.4%) specimens, with most specimens originating from the urban site (273) and the wet season (245). The mosquito species diversity consisted of *Culex pipiens pipiens* (280), *Culex* sp. (11), *C. neavei* (3), *C. nebulosus* (4), *C. bitaeniorhynchus* (1), *C. hutchinsoni* (1), *C. pipiens molestus* (1), *Aedes aegypti* (4), A. *simpsoni* (1), *Mansonia africana* (2), *Mansonia* sp. (1) and *Anopheles rivulorum* (1). A total of 35 molecular operational taxonomic units (MOTUs) were identified, belonging to approximately 30 genera, which we grouped into 11 generic host categories: human (Homininae, *Homo sapiens*), domestic mammal (*Felis catus*, *Canis sp.*, *Canis lupus familiaris*, *Bos spp.*, *Sus scrofa*, *Ovis sp.*), domestic bird (*Gallus gallus*, *Meleagris gallopavo*), non-human primate (*Chlorocebus sp.*), commensal rodent (*Mus musculus*, *Rattus spp.*, *Rattus norvegicus*), non-commensal rodent (*Acomys cahirinus*, *Mastomys natalensis*, *Meriones sp.*, *Lophuromys flavopunctatus*, *Zyzomys argurus*, *Praomys jacksoni*, Gerbillinae), shrew (*Crocidura spp.*, *Crocidura dsinezumi*, *Sylvisorex lunaris*), wild mammal (*Megaloglossus woermanni*, *Rhynchocyon sp.*), crow (Corvidae, *Corvus albus*), water bird (*Bubulcus ibis*, *Threskiornis aethiopicus*, *Fregata magnificens*), and other bird (*Philentoma pyrhoptera*, Passeroidea, *Colius striatus*, *Centropus superciliosus*).

Mosquitoes fed on an average of 1.35 ± 0.62 host groups (range: 1-4) and 1.36 ± 0.62 MOTUs (range: 1-4). At the genus level, the mean number of MOTUs used per mosquito was 1.20 ± 0.45 for *Aedes*, 1.0 for *Anopheles*, 1.36 ± 0.61 for *Culex*, and 2.0 ± 1.0 for *Mansonia*. Similar values were obtained for host groups averages (1.20 ± 0.45, 1.0, 1.35 ± 0.61, and 2.0 ± 1.0, respectively).

### - Host-feeding patterns

Due to the limited number of blood-fed or gravid *Aedes* (N=5), *Anopheles* (N=1), and *Mansonia* (N=3) specimens, no genus-specific host feeding patterns or matrices are presented for these taxa. Their feeding matrices are provided in the supplementary information (Mosquito_analysis). The feeding matrix of *Culex* (Figure 3) shows that the most often used hosts are humans (205/301; 68%), non-commensal rodents (67/301; 22%), and crows (55/301; 18%). The human category was detected significantly more frequent than all other host categories (p < 0.001), the same trend was observed for both non-commensal rodents and crows with the exception that they did not significantly differ in frequency from each other (χ²(df=1) = 1.1, p = 0.303) Figure 3. All other pairwise test results can be found in the appendix (Appendix A: McNemar Chi-square for *Culex*).

**Figure 3:**
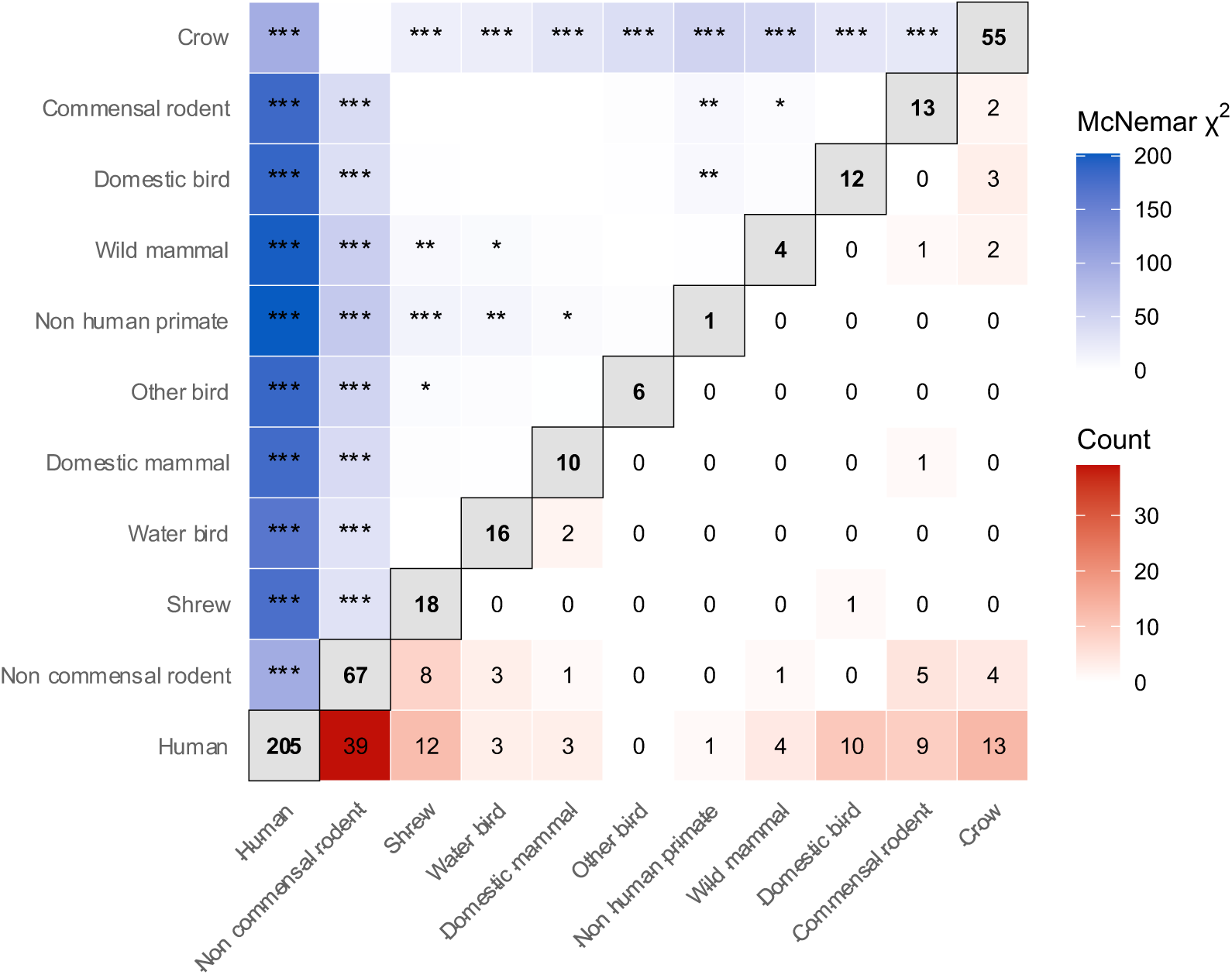
Feeding matrix of Culex specimens (N=301). The matrix visualizes host feeding overlap: diagonal values indicate how many mosquitoes that fed on each host, the lower triangle shows the number of mosquitoes that fed on both hosts, and the upper triangle shows the McNemar pairwise Chi-square (χ²) result in colour and the symbol represents the Holm adjusted p-value for multiple testing (* <0.05, ** <0.01, *** <0.001).

### - Human co-feedings

Of the 301 *Culex* specimens analysed, 205 had fed on humans. Host species significantly influenced human co-feeding patterns (GLMM, χ²(df8) = 88.16, p < 0.001), with human co-feeding on non-commensal rodents being more likely (Probability ± SE: 16.9% ± 3.0%) than on any other host (p ≤ 0.005) (Figure 4). All other pairwise test results can be found in the appendix (Appendix B: Human co-feeding pairwise results).

**Figure 4:**
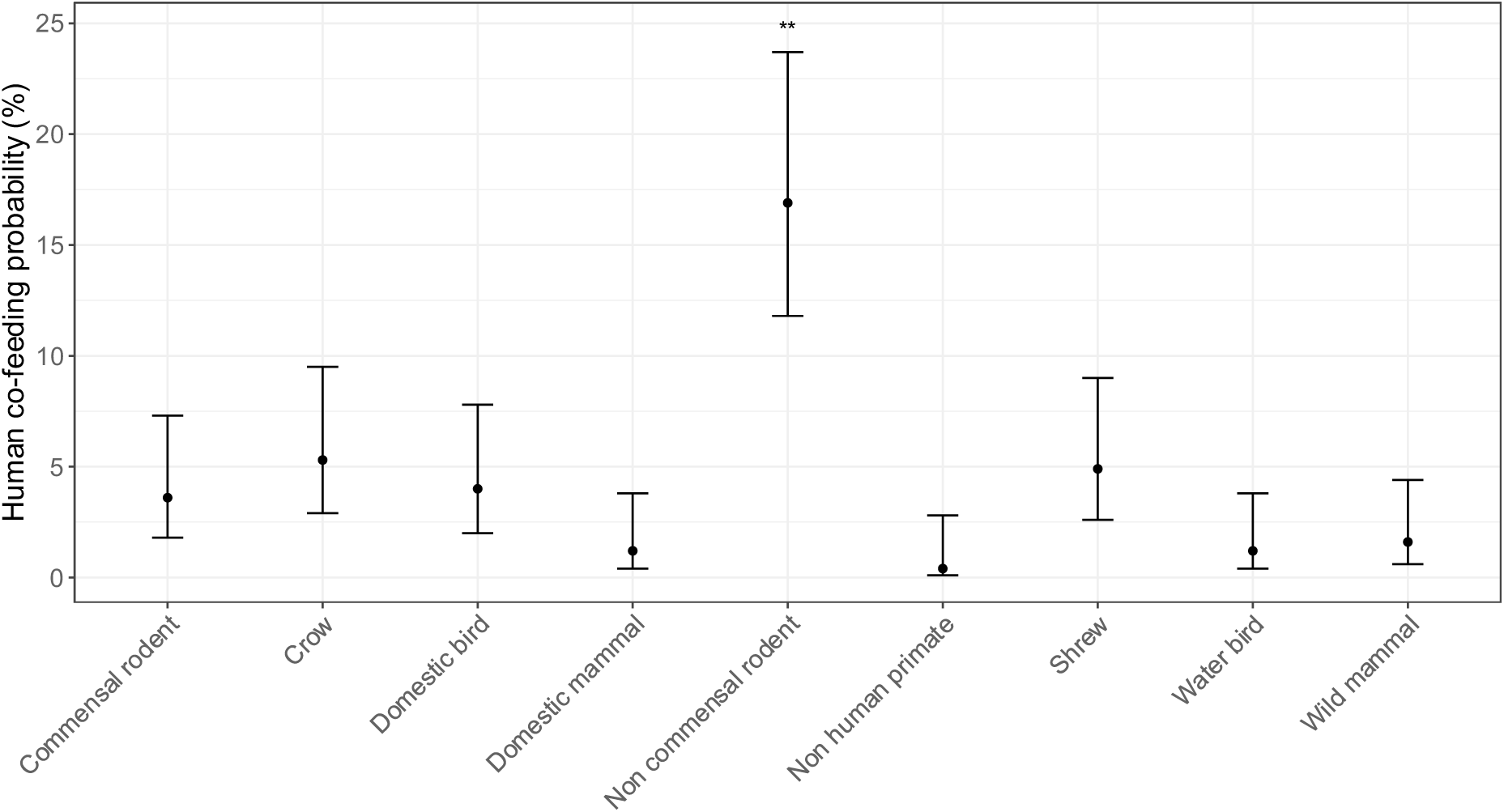
Human co-feeding (N=205) probability with a 95% confidence interval for each host group. The host group indicated with asterisks (**) means that it is significantly different from all other host groups (p<0.01).

### - Arbovirus detections

Among the 363 individual mosquito (i.e., blood-fed/gravid) and 292 pool extracts (i.e., non-blood-fed/non-gravid females) screened for arboviruses, three individual mosquitoes and 38 pools tested qPCR positive for *Orthoflavivirus*, while one pool tested qPCR positive for *Alphavirus*. Of the 297 samples tested specifically for WNV based on the at least 50% bird blood meal non were found positive. The six samples with more than 50% domestic mammal blood tested negative for RVFV. Sanger sequencing failed for 12 pools (11 *Orthoflavivirus*- and one *Alphavirus*-qPCR positive), whereas 26 *Orthoflavivirus*-positive pools yielded sequences of sufficient quality for alignment and further analysis. Nineteen samples, including the three blood-fed samples, clustered most closely with an endogenous viral element (EVE) in chromosome two of *Aedes aegypti*. Three samples clustered with each other with Culex flavivirus (NC_008604) being the nearest neighbour. Three other samples clustered basely in a clade of insect specific viruses. Two samples clustered with Bagaza virus (NC_012534) as their closest neighbour (Figure 5). See appendix for sample details and BLAST results (Appendix C: BLAST results table).

**Figure 5:**
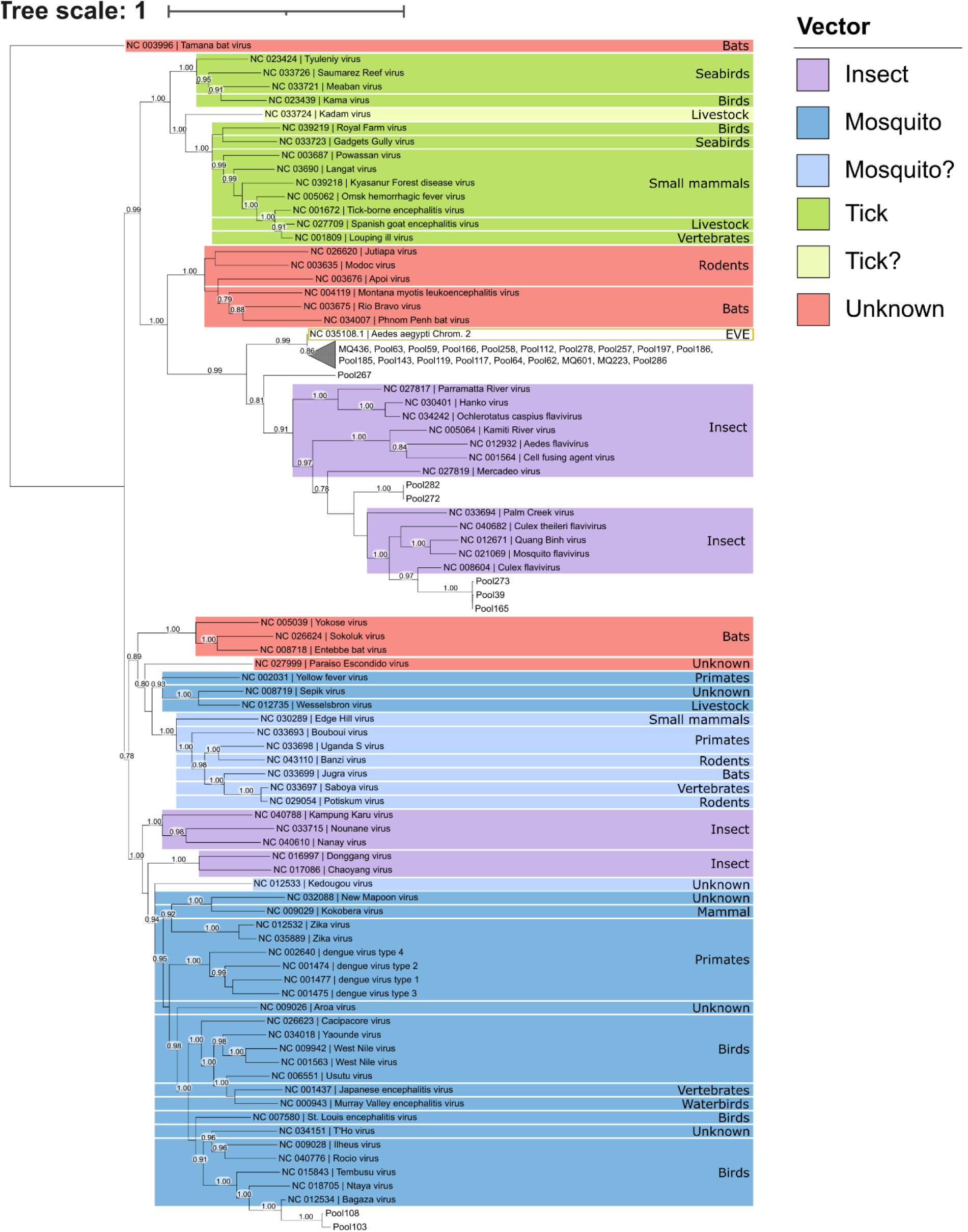
Orthoflavivirus non-structural protein 5 (NS5, ±2700bp) alignment for all known reference sequences, combined with the successful acquired amplicons (±230bp). The phylogenetic tree was created using FastTree 2.2 in Geneious version 2026.1 and visualized in ITOL [55]. Node numbers represent the support value above 75% and reference sequences are coloured by the known vector and list the known reservoir according to ICTV database [56]. The reference sequence of Tamana bat virus was used as the outgroup to root the tree [56].

## Discussion

This study aimed to identify the vertebrate hosts most frequently utilized by *Culex* mosquitoes and assess the extent to which they feed on multiple host groups. We hypothesized that mosquitoes feed preferentially on certain vertebrate species, and as such that the risk of arbovirus transmission from non-human hosts to humans is skewed towards originating from these preferred vertebrate species. Our findings generally supported this hypothesis, with *Culex* mosquitoes having a disproportionally feeding preference for humans, non-commensal rodents and crows, with co-feeding patterns demonstrating substantial host sharing between humans and non-commensal rodents.

The broad host use we detected, encompassing 35 MOTU’s, is consistent with previous studies demonstrating that *Culex* mosquitoes can exploit a wide range of vertebrate hosts and often display flexible feeding behaviour. In western Uganda, Crabtree et al. (2013) identified blood meals originating from humans, birds, rodents, bats, primates and a diversity of wild mammals [46]. The predominance of humans (68%) in our dataset contrasts with several African studies in which livestock represented the primary host group [24,47]. These differences likely reflect regional variation in host availability, habitat characteristics and mosquito community composition. Additionally, our sampling was performed in a highly urbanized environment and almost all blood fed specimens came from the city centre, where livestock is not housed. A recent global meta-analysis of *Culex* blood meals demonstrated that feeding patterns vary substantially among regions and that geographically distinct populations of the same species can show markedly different host associations [48].

The detection of multiple, on average 1.36 MOTUs within individual *Culex* mosquitoes suggests that host switching and co-feeding is not uncommon. Likewise, other studies found that East African culicine mosquitoes frequently fed on multiple host species within the same gonotrophic cycle [24,47,49]. This co-feeding behaviour may facilitate pathogen movement between otherwise disconnected host groups, creating opportunities for arbovirus spillover between wildlife and humans, potentially linking sylvatic and urban transmission cycles.

A notable finding was the frequent occurrence of non-commensal rodent blood meals (22%) and their strong association with the co-occurrence of human blood meals (p ≤ 0.003). Rodents are relatively underrepresented in the East African mosquito blood-meal literature. Several studies from the region primarily focused on mosquitoes feeding on humans, livestock, and domestic animals. For example, Muturi et al. (2008) and Munyao et al. (2020) used direct enzyme-linked immunosorbent assays (ELISA) targeting a predefined set of common vertebrate hosts that did not include rodents, despite acknowledging their presence in the study area and role in mosquito blood feeding use [24,47]. Likewise, the high-resolution melting analyses applied by Musa et al. (2020) focused mainly on humans, livestock, domestic animals, and wild ungulates, without incorporating rodent reference profiles for host identification [49].

Consequently, the contribution of rodents to mosquito feeding patterns may have been underestimated in these studies, especially since rodents have been detected when broader host identification approaches were applied. Omondi et al. (2015) identified both rodent blood meals and rodent-human co-feeding events at wildlife-livestock interfaces, while Tchouassi et al. (2024) detected rodents as both single-host blood meals and components of mixed blood meals involving humans and birds [23,25]. Similarly, Crabtree et al. (2013) reported rodent-derived blood meals in several mosquito species [46]. Although these studies demonstrate that rodents are incorporated into mosquito feeding networks in East Africa, they generally constituted only a minor proportion of identified blood meals and were not considered major host groups.

In contrast, rodents accounted for a substantial proportion of blood meals in our study, suggesting they may occupy a more prominent role in local mosquito-host interaction networks than previously appreciated. This difference may partly reflect methodological advances. By applying nanopore-based next-generation sequencing, we were able to identify a broad range of vertebrate molecular taxonomic units without restricting analyses to a predefined set of potential host species. This approach additionally enabled discrimination among multiple rodent taxa, providing a more detailed characterization of mosquito-rodent interactions. Ecological differences between study systems may also have contributed to the observed pattern. Our sampling was conducted in a highly urbanized environment where vertebrate diversity is comparatively limited and where rodents likely represent one of the most abundant non-human mammalian groups. Under such conditions, rodents may serve as one of the few readily available alternative hosts for mosquitoes, potentially explaining both the high prevalence of rodent blood meals and their frequent co-occurrence with human blood meals. Together, these findings suggest that rodents may play a more important and potentially preferential role in local mosquito feeding ecology than indicated by previous studies from the region. The role of rodents in mosquito feeding networks may also have epidemiological implications. Previous studies have reported viral RNA, virus isolation, or arbovirus-specific antibodies in rodent populations, suggesting exposure to medically important arboviruses. For example, Usutu virus has been isolated from rodents in Senegal, while antibodies against several arboviruses, including dengue virus, chikungunya virus, and West Nile virus, have been detected in non-commensal rodent populations from Tanzania [51,52]. Although the competence of rodents for arbovirus transmission remains unclear, these observations indicate that rodents interact with arboviruses in natural systems. When considered alongside the high frequency of rodent-derived blood meals observed in the present study, these findings support the hypothesis that rodents may contribute to local arbovirus transmission cycles and warrant further investigation as potential reservoir or amplification hosts.

Despite the extensive host sharing observed in this study, screening of mosquitoes did not reveal any human-pathogenic arboviruses. Instead, positive detections were limited to insect-specific viruses, an endogenous viral element in chromosome 2 of *Aedes aegypti*, and two detections of Bagaza virus, an emerging *Orthoflavivirus* frequently associated with disease and mortality in wild birds [50]. The absence of human-pathogenic arboviruses is consistent with numerous vector surveillance studies in which active arboviral circulation is only detected sporadically or not all [51–54]. The lack of human-pathogenic arboviruses in our dataset may therefore reflect low infection prevalence, temporal variation in arbovirus circulation, or limitations inherent to snapshot surveillance rather than the absence of transmission risk. Nevertheless, the detection of Bagaza virus demonstrates that arboviruses are circulating within the local mosquito community and highlights the value of integrated virus screening and blood-meal analysis for identifying potential transmission networks.

Several limitations should be considered when interpreting our results. The 12S barcoding fragment did not always provide sufficient taxonomic resolution for species-level host identification, resulting in some blood meals being assigned only to the genus or family level. Consequently, some host diversity remains unresolved due to an incomplete 12S reference database for all known vertebrates. In addition, arbovirus screening mainly relied on broad-range genus-level qPCR assays, which prioritize broad detection over specificity. Furthermore, SYBR Green-based assays generally offer lower specificity than probe-based approaches. The approach used here successfully enabled the simultaneous characterization of vertebrate blood-meal sources and arbovirus screening within mosquito populations. Expanding the marker set in future applications would further increase host identification accuracy and taxonomic coverage, providing an even more powerful tool for studies of vector-host-pathogen networks.

## Conclusion

Our study demonstrates that *Culex* mosquitoes exhibit broad and flexible host use, feeding on at least 35 MOTUs and frequently sharing blood meals among different taxonomic units. Humans were the preferential blood-meal source, but the frequent detection of multiple host species belonging to distinct ecological settings within individual mosquitoes highlights the potential for *Culex* mosquitoes to act as bridge vectors, facilitating pathogen movement across host communities.

A key finding was the significant associations of non-commensal rodents with human co-feeding, suggesting that rodents are also a preferred blood host and may contribute more substantially to local arbovirus transmission dynamics than previously recognized and should be considered more explicitly in future surveillance and ecological studies.

Although no human-pathogenic arboviruses were detected, the identification of bird-pathogenic Bagaza virus and insect-specific viral sequences indicates detectable *Orthoflavivirus* circulation within the local mosquito community. Overall, this study highlights the value of integrating blood-meal analysis and virus screening to improve our understanding of mosquito-host interactions and potential pathways of arbovirus transmission and spillover.

## Supporting information

Nanopore_SamplePrep

Bioinformatics

Mosquito analysis

## Acknowledgements

We would like to thank Geoffrey Sabuni, Shabani Lutea and Heleen Bulcke for their assistance during the fieldwork.

## Appendix

**Appendix A:** Pairwise McNemar chi-square tests comparing the occurrence of host categories within blood meals of *Culex* mosquitoes. Chi-square statistics with Holm-adjusted *p*-values to account for multiple testing and one degree of freedom.

| Host category 1 | Host category 2 | Chi-square | p-value |
|---|---|---|---|
| **Human** | **Non commensal rodent** | **96.75** | **<0.001** |
| **Human** | **Shrew** | **173.85** | **<0.001** |
| **Human** | **Water bird** | **164.39** | **<0.001** |
| **Human** | **Domestic mammal** | **180.08** | **<0.001** |
| **Human** | **Other bird** | **185.80** | **<0.001** |
| **Human** | **Non human primate** | **202.00** | **<0.001** |
| **Human** | **Wild mammal** | **199.00** | **<0.001** |
| **Human** | **Domestic bird** | **187.13** | **<0.001** |
| **Human** | **Commensal rodent** | **182.40** | **<0.001** |
| **Human** | **Crow** | **94.88** | **<0.001** |
| **Non commensal rodent** | **Shrew** | **33.39** | **<0.001** |
| **Non commensal rodent** | **Water bird** | **32.47** | **<0.001** |
| **Non commensal rodent** | **Domestic mammal** | **41.81** | **<0.001** |
| **Non commensal rodent** | **Other bird** | **49.32** | **<0.001** |
| **Non commensal rodent** | **Non human primate** | **62.13** | **<0.001** |
| **Non commensal rodent** | **Wild mammal** | **55.71** | **<0.001** |
| **Non commensal rodent** | **Domestic bird** | **36.91** | **<0.001** |
| **Non commensal rodent** | **Commensal rodent** | **40.13** | **<0.001** |
| Non commensal rodent | Crow | 1.06 | 0.303 |
| Shrew | Water bird | 0.03 | 0.864 |
| Shrew | Domestic mammal | 1.75 | 0.186 |
| **Shrew** | **Other bird** | **5.04** | **0.025** |
| **Shrew** | **Non human primate** | **13.47** | **<0.001** |
| **Shrew** | **Wild mammal** | **7.68** | **0.006** |
| Shrew | Domestic bird | 0.89 | 0.345 |
| Shrew | Commensal rodent | 0.52 | 0.472 |
| **Shrew** | **Crow** | **17.75** | **<0.001** |
| Water bird | Domestic mammal | 1.14 | 0.286 |
| Water bird | Other bird | 3.68 | 0.055 |
| **Water bird** | **Non human primate** | **11.53** | **0.001** |
| **Water bird** | **Wild mammal** | **6.05** | **0.014** |
| Water bird | Domestic bird | 0.32 | 0.571 |
| Water bird | Commensal rodent | 0.14 | 0.710 |
| **Water bird** | **Crow** | **20.34** | **<0.001** |
| Domestic mammal | Other bird | 0.56 | 0.453 |
| **Domestic mammal** | **Non human primate** | **5.82** | **0.016** |
| Domestic mammal | Wild mammal | 1.79 | 0.181 |
| Domestic mammal | Domestic bird | 0.05 | 0.831 |
| Domestic mammal | Commensal rodent | 0.19 | 0.663 |
| **Domestic mammal** | **Crow** | **29.78** | **<0.001** |
| Other bird | Non human primate | 2.29 | 0.131 |
| Other bird | Wild mammal | 0.10 | 0.752 |
| Other bird | Domestic bird | 1.39 | 0.239 |
| Other bird | Commensal rodent | 1.89 | 0.169 |
| **Other bird** | **Crow** | **37.77** | **<0.001** |
| Non human primate | Wild mammal | 0.80 | 0.371 |
| **Non human primate** | **Domestic bird** | **7.69** | **0.006** |
| **Non human primate** | **Commensal rodent** | **8.64** | **0.003** |
| **Non human primate** | **Crow** | **50.16** | **<0.001** |
| Wild mammal | Domestic bird | 3.06 | 0.080 |
| **Wild mammal** | **Commensal rodent** | **4.27** | **0.039** |
| **Wild mammal** | **Crow** | **45.45** | **<0.001** |
| Domestic bird | Commensal rodent | 0.00 | 1.000 |
| **Domestic bird** | **Crow** | **28.92** | **<0.001** |
| **Commensal rodent** | **Crow** | **26.27** | **<0.001** |

**Appendix B:** Pairwise comparisons human co-feeding probability among host categories. Comparisons were performed using estimated marginal means derived from a generalized linear mixed-effects model (GLMM) with a binomial error distribution. Values represent odds ratios between host categories, with Tukey-adjusted *p*-values to account for multiple comparisons. Odds ratios >1 indicate that the first host category was more likely to co-occur than the second host category, whereas odds ratios <1 indicate the opposite. Significant comparisons (*p* < 0.05) are highlighted in bold.

| Comparison | Odds ratio | z-ratio | Adjusted p-value |
|---|---|---|---|
| Commensal rodent / Crow | 0.674 | -0.879 | 0.994 |
| Commensal rodent / Domestic bird | 0.895 | -0.234 | 1.000 |
| Commensal rodent / Domestic mammal | 3.120 | 1.682 | 0.758 |
| **Commensal rodent / Non commensal rodent** | **0.184** | **-4.316** | **<0.001** |
| Commensal rodent / Non human primate | 9.519 | 2.124 | 0.457 |
| Commensal rodent / Shrew | 0.734 | -0.676 | 0.999 |
| Commensal rodent / Water bird | 3.142 | 1.688 | 0.754 |
| Commensal rodent / Wild mammal | 2.323 | 1.378 | 0.906 |
| Crow / Domestic bird | 1.329 | 0.650 | 0.999 |
| Crow / Domestic mammal | 4.631 | 2.355 | 0.309 |
| **Crow / Non commensal rodent** | **0.273** | **-3.756** | **0.005** |
| Crow / Non human primate | 14.130 | 2.535 | 0.215 |
| Crow / Shrew | 1.090 | 0.207 | 1.000 |
| Crow / Water bird | 4.664 | 2.359 | 0.307 |
| Crow / Wild mammal | 3.449 | 2.122 | 0.458 |
| Domestic bird / Domestic mammal | 3.485 | 1.868 | 0.636 |
| **Domestic bird / Non** | **0.206** | **-4.187** | **<0.001** |
| Domestic bird / Non human primate | 10.634 | 2.239 | 0.380 |
| Domestic bird / Shrew | 0.821 | -0.445 | 1.000 |
| Domestic bird / Water bird | 3.510 | 1.874 | 0.632 |
| Domestic bird / Wild mammal | 2.596 | 1.582 | 0.815 |
| **Domestic mammal / Non commensal rodent** | **0.059** | **-4.616** | **<0.001** |
| Domestic mammal / Non human primate | 3.051 | 0.961 | 0.989 |
| Domestic mammal / Shrew | 0.235 | -2.206 | 0.402 |
| Domestic mammal / Water bird | 1.007 | 0.009 | 1.000 |
| Domestic mammal / Wild mammal | 0.745 | -0.382 | 1.000 |
| **Non commensal rodent / Non human primate** | **51.674** | **3.861** | **0.004** |
| **Non commensal rodent /Shrew** | **3.987** | **3.902** | **0.003** |
| **Non commensal rodent / Water bird** | **17.057** | **4.614** | **<0.001** |
| **Non commensal rodent / Wild mammal** | **12.613** | **4.688** | **<0.001** |
| Non human primate / Shrew | 0.077 | -2.445 | 0.260 |
| Non human primate / Water bird | 0.330 | -0.954 | 0.990 |
| Non human primate / Wild mammal | 0.244 | -1.255 | 0.944 |
| Shrew / Water bird | 4.278 | 2.211 | 0.398 |
| Shrew / Wild mammal | 3.164 | 1.956 | 0.574 |
| Water bird / Wild mammal | 0.739 | -0.390 | 1.000 |

**Appendix C:** Blast results table. Successful sequences that were compared using BLASTn to all known Orthoflavivirus reference sequences from NCBI. Sample ID, pool size, identification of the mosquito species and site are included, as well as the results from the BLASTn comparison (matched reference sequence, query cover and pairwise identity).

| Sample | Pool size | Sample species | Site | Match | Query cover (%) | Pairwise Id. (%) | Ambiguities | Sequence |
|---|---|---|---|---|---|---|---|---|
| MQ223 | 1 | *Aedes aegypti* | Urban | Chrom2 of *Aedes aegypti* (NC_035108.1) | 100 | 99.6 | 0 | Assembly |
| MQ601 | 1 | *Aedes aegypti* | urban | Chrom2 of *Aedes aegypti* (NC_035108.1) | 100 | 99.6 | 0 | Assembly |
| MQ436 | 1 | *Aedes aegypti* | Suburban | Chrom2 of *Aedes aegypti* (NC_035108.1) | 100 | 99.0 | 2 | Assembly |
| Pool185 | 17 | *Aedes spp.* | Urban | Chrom2 of *Aedes aegypti* (NC_035108.1) | 100 | 99.6 | 0 | Assembly |
| Pool117 | 16 | *Aedes spp.* | Suburban | Chrom2 of *Aedes aegypti* (NC_035108.1) | 100 | 99.6 | 0 | Assembly |
| Pool119 | 16 | *Aedes spp.* | Suburban | Chrom2 of *Aedes aegypti* (NC_035108.1) | 100 | 99.6 | 0 | Assembly |
| Pool143 | 16 | *Aedes spp.* | Suburban | Chrom2 of *Aedes aegypti* (NC_035108.1) | 100 | 99.6 | 0 | Assembly |
| Pool197 | 16 | *Aedes spp.* | Suburban | Chrom2 of *Aedes aegypti* (NC_035108.1) | 100 | 99.6 | 0 | Assembly |
| Pool278 | 12 | *Aedes spp.* | Suburban | Chrom2 of *Aedes aegypti* (NC_035108.1) | 100 | 99.6 | 0 | Assembly |
| Pool286 | 12 | *Aedes spp.* | Suburban | Chrom2 of *Aedes aegypti* (NC_035108.1) | 100 | 99.6 | 0 | Assembly |
| Pool64 | 16 | *Aedes spp.* | Suburban | Chrom2 of *Aedes aegypti* (NC_035108.1) | 100 | 99.6 | 0 | Assembly |
| Pool258 | 12 | *Aedes spp.* | Suburban | Chrom2 of *Aedes aegypti* (NC_035108.1) | 100 | 99.3 | 1 | Assembly |
| Pool62 | 16 | *Aedes spp.* | Rural | Chrom2 of *Aedes aegypti* (NC_035108.1) | 100 | 99.6 | 0 | Assembly |
| Pool186 | 16 | *Aedes spp.* | Rural | Chrom2 of *Aedes aegypti* (NC_035108.1) | 100 | 99.6 | 0 | Assembly |
| Pool257 | 12 | *Aedes spp.* | Rural | Chrom2 of *Aedes aegypti* (NC_035108.1) | 100 | 99.6 | 0 | Assembly |
| Pool112 | 16 | *Aedes spp.* | Rural | Chrom2 of *Aedes aegypti* (NC_035108.1) | 100 | 99.3 | 1 | Assembly |
| Pool63 | 17 | *Aedes spp.* | Rural | Chrom2 of *Aedes aegypti* (NC_035108.1) | 100 | 98.5 | 0 | Assembly |
| Pool165 | 16 | *Culex spp.* | Urban | Culex flavivirus (NC_008604) | 100 | 93.0 | 0 | Assembly |
| Pool39 | 16 | *Culex spp.* | Urban | Culex flavivirus (NC_008604) | 100 | 92.6 | 0 | Assembly |
| Pool103 | 16 | *Culex spp.* | Rural | *Bagaza virus* (NC_012534) | 100 | 94.4 | 0 | Assembly |
| Pool108 | 16 | *Culex spp.* | Rural | *Bagaza virus* (NC_012534) | 100 | 93.1 | 1 | Assembly |
| Pool273 | 14 | *Culex spp.* | Rural | *Culex flavivirus* (NC_008604) | 100 | 92.60 | 0 | Assembly |
| Pool267 | 2 | *Mansonia spp.* | Suburban | *Culex theileri flavivirus* (NC_040682) | 91.70 | 77.10 | 0 | Assembly |
| Pool282 | 14 | *Mansonia spp.* | Rural | *Culex theileri flavivirus* (NC_040682) | 98.29 | 81.2 | 1 | Assembly |
| Pool272 | 14 | *Mansonia spp.* | Rural | *Culex theileri flavivirus* (NC_040682) | 100 | 84.1 | 1 | Reverse |
| Pool59 | 16 | Unknown | Rural | Chrom2 of *Aedes aegypti* (NC_035108.1) | 100 | 99.5 | 0 | Assembly |
| Pool166 | 16 | *Culex spp.* | Urban | Chrom2 of *Aedes aegypti* (NC_035108.1) | 100 | 99.0 | 0 | Reverse |

## Supplementary information

Nanopore_SamplePrep.pdf

Bioinformatics.html

Mosquito_analysis.html

