## Supplementary material for "Blood-meal analysis indicates preferential feeding on humans and non-commensal rodents by *Culex* mosquitoes in Tanzania": Nanopore_SamplePrep

### Nanopore protocols

#### Condensed version (WimDK)

A. PCR
B. PCR clean-up
C. END-prep DNA
D. Bead purification
E. Native barcode ligation
F. Adapter ligation
G. Priming and loading
H. Flow cell wash
I. Store flow cell

##### References

Ligation sequencing amplicons - Native Barcoding Kit 96 V14 (SQK-NBD114.96)
https://nanoporetech.com/document/ligation-sequencing-amplicons-native-barcoding-v14-sqk-nbd114-96
Flow Cell Wash Kit (EXP-WSH004 or EXP-WSH004-XL)
https://nanoporetech.com/document/flow-cell-wash-kit-exp-wsh004#overview-of-the-protocol

---

#### A. PCR

Perform your desired PCR
(Measure the COX1 concentration with Qubit)

#### B. PCR clean-up

(merge 12S amplicons and 100ng of COI amplicon)

*(Macherey-Nagel: NucleoSpin® Gel and PCR Clean-up; page 17; 5 Protocols)*

Before starting the preparation:

- Check if Wash Buffer NT3 was prepared according to section 3.

---

1. Adjust DNA binding condition

*For very small sample volumes < 30 µL adjust the volume of the reaction mixture to 50–100 µL with water.*

Mix **1 volume of sample** with **2 volumes** of **Buffer NTI** (e.g., mix 50 µL PCR reaction and 100 µL Buffer NTI).

*Mix should be green, if yellow add more NTI (titration).*

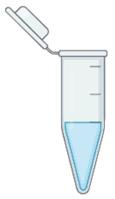

+ 2 vol. NTI per 1 vol. sample

vol. sample >50µL

---

2. Bind DNA

Place a **NucleoSpin® Gel and PCR Clean-up Column** into a Collection Tube (2 mL) and load up to 700 µL sample. Centrifuge for **30 s** at **11,000 x g**. Discard flow-through and place the column back into the collection tube.

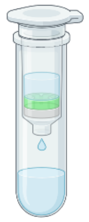

Load sample

↻ 11,000 x g 30 s

---

3. Wash silica membrane

Add **700 µL Buffer NT3** to the NucleoSpin® Gel and PCR Clean-up Column. Centrifuge for **30 s** at **11,000 x g**. Discard flow-through and place the column back into the collection tube.

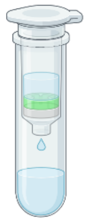

+ 700 µL NT3

↻ 11,000 x g 30 s

---

4. Dry silica membrane

Centrifuge for **1 min** at **11,000 x g** to remove Buffer NT3 completely.

*Make sure the spin column does not come in contact with the flow-through while removing it from the centrifuge and the collection tube.*

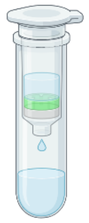

↻ 11,000 x g 1 min

---

5. Elute DNA

Place the NucleoSpin® Gel and PCR Clean-up Column into a new 1.5 mL microcentrifuge tube (not provided).

Add **30 µL Buffer NE** and incubate at **room temperature** (15–25 °C) for **1 min**. Centrifuge for **1 min** at **11,000 x g**.

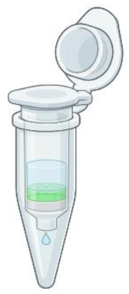

+ 30 µL NE

Room temp. 1 min

↻ 11,000 x g 1 min

#### C. <u>END-prep DNA</u>

1. Thaw DNA control Sample (DCS) at room temperature, vortex and keep on ice
2. Prepare the NEBNext Ultra II End Repair / dA-tailing Module reagents
   2.1. Thaw on ice or fridge
   2.2. Mix reagents (**do not vortex Ultra II End Prep Enzyme Mix**)
      2.2.1. NEBNext Ultra II End Prep Reaction Buffer is mixed by **vortex**
   2.3. Spin down reagents
3. Dilute DCS
   3.1. Add 105 µL elution buffer (EB) to one DCS tube
   3.2. Mix by pipetting and spin down
   3.3. One tube is for 140 samples, store at -20°C
4. PCR tubes with **11.5 µL of DNA sample** (if omitting DCS, take 12.5 µL)
5. Add following master mix, **3.5 µL per sample**

| *Reagents* | *Volume/sample* | *24 samples (2 extra)* | *16 samples (2 extra)* |
|---|---|---|---|
| *DCS* | 1 µL | **26 µL** | **18 µL** |
| *Ultra II End-prep Reaction Buffer* | 1.75 µL | **45.5 µL** | **31.5 µL** |
| *Ultra II End-prep Enzyme Mix* | 0.75 µL | **19.5µL** | **13.5 µL** |
| *Master mix* | 3.5 µL | 91 µL | 63 µL |
| *DNA sample* | 11.5 µL | | |
| *Total* | 15 µL | | |

6. Mix by pipetting, spin down
7. Thermal cycler (*PCR 3 program END0prep)*, incubate at **20°C for 5 minutes** and **65°C for 5 minutes**. Put on ice afterwards, and spin down. (Optional Qubit measurement)

**Perform the BEAD PURIFICATION for storage of the END-prepped samples!**
**If no storage, skip bead purification and continue with native barcode ligation!**

#### D. Bead purification

(extra AMPure beads required)

Purifying a sample to obtain pure DNA used for subsequent reaction.

*If not already, transfer the sample to a clean 1.5 mL Eppendorf tube (LoBind) or a clean 200 µL PCR tube/strip.*

*e.g. 15µL sample*

1. Resuspend the AMPure XP beads stock by vortexing. **Take an aliquot** of the bead solution to avoid stock contamination.
2. Add 1.8X of AMPure XP beads in relation to sample volume, flick mix (**DO NOT VORTEX**), pulse centrifuge 2-3 seconds (**no excessive centrifugation**).

   e.g. **27 µL AMPure XP beads to 15 µL sample**
3. Incubate at room temperature for 5 minutes (avoid excess incubation)
4. Open lids and place tubes in magnetic rack.
5. Wait until elute is clear (± 2 min) and pellet has formed against the wall.
6. Aspirate the liquid gently (**do not disturb the bead pellet**).
7. **Add 200 µL of 90% ethanol** on the opposite side of the bead pellet, wait 30 seconds and aspirate the ethanol.
8. Remove from magnet and pulse centrifuge
9. Open lids gently and put tubes on magnetic rack.
10. Wait until beads form a pellet and aspirate remaining ethanol with a P10 pipet, dry pellet for 30 seconds (**no cracks**)
11. **Add 15 µL NFW** on the side of the pellet (the liquid should run over the pellet)
12. Remove from magnet, flick mix and pulse centrifuge. **Incubate** the tubes for **2-3 minutes** at **room temperature** (**not in the magnet**).
13. Open lids gently and put tubes on magnetic rack. Wait until bead pellet is formed, **aspirate NFW containing DNA** and **transfer to a new clean tube.**

#### E. Native barcode ligation

1. Thaw NEB Blunt/TA Ligase Master Mix, AMPure XP Beads, EDTA, Native barcodes (NB01-96; keep on ice) at room temperature, flick mix and spin down
2. Measure Qubit for each sample, adjust concentrations to have **equimolar DNA**
3. In a clean 96-well plate add reagents **in following order** per well and mix by pipetting after each addition, spin down after completion of plate

| *Reagent* | *Volume (µL)* |
|---|---|
| *End-prepped DNA* | **3.75** |
| *Native Barcode (NB01-96)* | **1.25** |
| *Blunt/TA Ligase Master Mix* | **5** |

4. Thermal cycler (*PCR 3 program BarcodeLG*), incubate at **room temperature** (18-23°C) **for 20 min**. Put on ice afterwards, and spin down.
5. Add EDTA to each well, mix by pipetting and briefly spin down
   5.1. **Clear cap EDTA 1 µL** or **Blue cap EDTA 2 µL**
6. Pool all barcoded samples in a 1.5 mL Eppendorf DNA LoBind tube. Check how many samples! #Samples **>60 option A** otherwise (**<60) option B**

| A. Pool in **two tubes** and repeat next steps for each tube! | B. Pool in **one** tube |
|---|---|

7. Resuspend AMPure XP Beads by vortex, add 1X AMPure XP Beads to the pooled barcode tube, mix by pipetting and flick mix, pulse centrifuge (no excessive centrifugation) (BeadVol(µL) = #Samples*11 or #Samples*12)

| A. BeadVol ÷ 2 = ........ µL Beads (per tube) | B. ........ µL Beads |
|---|---|

8. Incubate for 10 min at room temperature (18-23°C)
9. Spin down, open lid and place on magnetic rack
10. Wait until elute is clear and bead pellet has formed against the wall
11. Aspirate liquid (**do not disturb bead pellet**)
12. Add **700 µL SFB**, flick mix, spin down, return to magnet wait until elute is clear and bead pellet has formed, aspirate liquid
13. Repeat previous step
14. Remove from magnet, spin down, place back on magnet, open lid
15. Wait until beads form a pellet and aspirate the remaining liquid with a P10 pipet, dry pellet for 30 seconds (**no cracks**)
16. Remove from magnet, resuspend pellet in NFW, flick mix

| A. 17.5 µL per tube | B. 35 µL |
|---|---|

17. Incubate for 10 min at 37°C, flick mix every 2 min
18. Place on magnet wait until eluate is clear and bead pellet has formed
19. **Aspirate eluate** to a new 1.5 mL Eppendorf DNA LoBind tube (Pooled DNA sample!)

| A. Combine both eluates |
|---|

20. Take Qubit measurement, safe storage at 4°C overnight

#### F. Adapter ligation

1. Thaw NEBNext Quick Ligation Reaction Buffer at room temperature, flick mix and spin down
2. Thaw Native Adapter (NA) and Quick T4 DNA Ligase at room temperature, flick mix, spin down and place on ice
   2.1. **Do not vortex Quick T4 DNA Ligase**
3. Thaw Elution Buffer (EB) at room temperature, vortex and spin down
4. Thaw Short Fragment Buffer (SFB) at room temperature, vortex and spin down
5. In 1.5mL Eppendorf LoBind tube mix in **following order**, flick mix and spin down

| *Reagent* | *Volume (µL)* |
|---|---|
| Pooled barcoded sample | **30** |
| Native Adapter (NA) | **5** |
| NEBNext Quick Ligation Reaction Buffer (5X) | **10** |
| Quick T4 DNA Ligase | **5** |
| *Total* | 50 |

6. Incubate for **20 min at room temperature** (18-23°C)
7. Resuspend AMPure XP Beads by vortex, add 1X AMPure XP Beads
   7.1. Add **50 µL of AMPure XP Beads**, flick mix, spin down
8. Incubate for **10 minutes** at room temperature (18-23°C)
9. Spin down, put on magnet, open lid, wait until elute is clear and pellet is formed, aspirate liquid (**do not disturb pellet**)
10. Wash the beads with **125 µL of SFB**, remove from magnet, flick mix and spin down
11. Put on magnet, open lid, wait until elute is clear and pellet is formed, aspirate liquid
12. Repeat previous step
13. Remove tube from magnet and resuspend beads in Elution Buffer (EB)
    13.1. Add **15 µL Elution Buffer (EB)**, flick mix, spin down
14. Incubate for 10 min at 37°C, flick mix every 2 min
15. Place on magnet wait until eluate is clear and bead pellet has formed (1 min)
16. Extract the liquid from the tube and transfer to a new clean tube
17. Take Qubit measurement, safe storage at 4°C overnight
    17.1. Aim is 30-50 ng/µL

### G. Priming and loading

1. Take flow cell out of fridge, let acclimate to room temperature for 20 min (only R10.4.1 flow cells, FLO-MIN114)

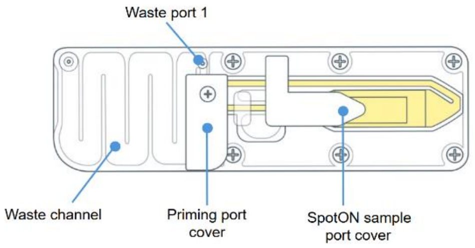


2. Thaw Sequencing Buffer (SB), Library Beads (LIB), Flow Cell Tether (FCT) and Flow Cell Flush (FCF) at room temperature. Vortex, spin down, keep on ice.

3. In 1.5mL Eppendorf LoBind tube make flow cell priming mix, mix by inverting and pipet mix at room temperature

| *Reagent* | *Volume (µL)* |
|---|---|
| Flow Cell Flush (FCF) | **1170** |
| Flow Cell Tether (FCT) | **30** |
| *Final volume* | 1200 |

4. Open MinION device, slide flow cell under the clip. Press down on priming port cover, ensure correct contact

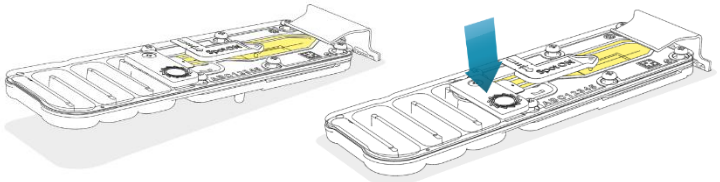

5. Connect MinION to laptop, run flow cell check for sufficient available pores (**NOT AFTER FLOW CELL WASH!**)
6. Slide priming port cover clockwise to access priming port

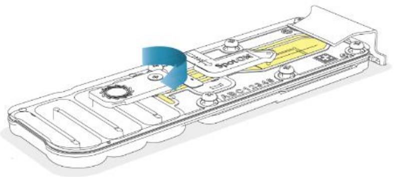

7. Check for air bubble, draw back **max 20-30 µL** (or until you see liquid in pipet tip)
   7.1. Set P1000 to 200µL, insert tip into priming port, turn **pipet volume up to max 220-230µL**, or until volume in tip

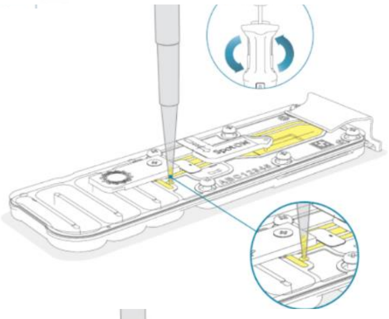

8. Load **800 µL of priming mix** in priming port (>10 sec)
   8.1. P1000 to 1000 µL take priming mix (**NO AIR BUBBLES!**)
   8.2. Insert tip into priming port, turn pipet volume down to 200µL by twisting pipet wheel slowly and continuous
   8.3. Close priming port cover and incubate for 5 minutes
   8.4. If after flow cell wash, remove waste buffers through waste port, P1000 to 1000 µL

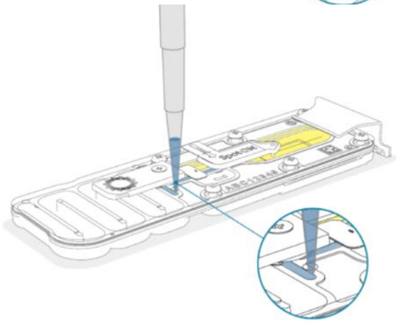

9. In 1.5mL Eppendorf LoBind tube make the final library, thoroughly mix the Library Beads (LIB) by pipetting before adding

| *Reagent* | *Volume (µL)* |
|---|---|
| Sequencing Buffer (SB) | **37.5** |
| Library Beads (LIB) | **25.5** |
| DNA library | **12** (30-50 ng/µL) |
| *Total* | 75 |

10. Open SpotON sample port and priming port cover. Load **200µL of priming mix** into flow cell priming port (**NOT SpotOn!**) P1000 to 400 µL, turn wheel slow and steady, **NO AIR BUBBLES!**, >5 sec

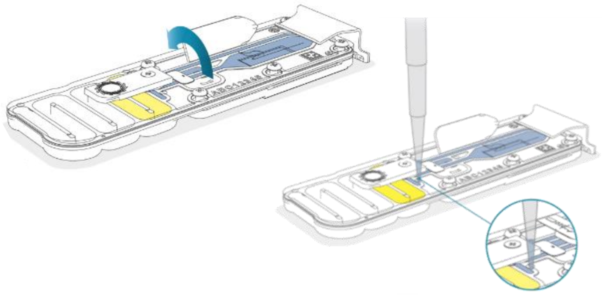

11. Mix the final library by gently pipetting prior to loading
12. Load **75 µL of final library** in the SpotON sample port in a dropwise fashion, ensure each drop flows in the port before the next one

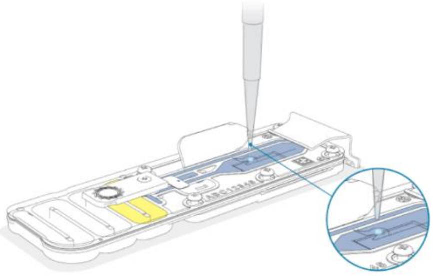

13. Gently close SpotON sample port cover, check bung enters the SpotON port. Close the priming port

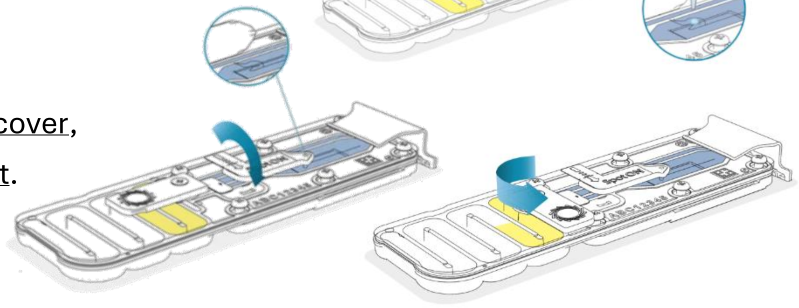

14. Place light shield on flow cell, place against the clip (**do not force under clip**)

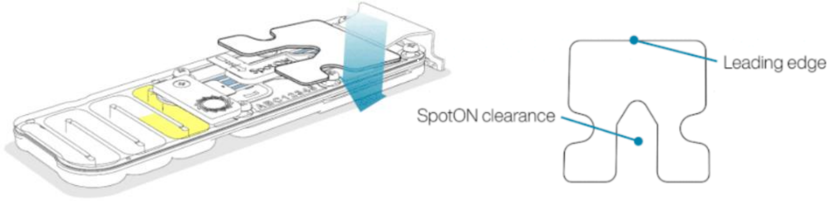


15. Close MinION and set up sequencing run on MinKNOW
    15.1. SQK-NBD114.96

### H. Flow cell wash

1. Thaw Wash Mix (WMX) on ice, do not vortex. Thaw Wash Diluent (DIL) at room temperature and vortex, spin down and keep on ice
2. In 1.5mL Eppendorf LoBind tube make the Flow Cell Wash Mix, mix by pipetting, do not vortex, keep on ice.

| ***Reagent*** | **Volume (µL)** |
|---|---|
| *Wash Mix (WMX)* | **2** |
| *Wash Diluent (DIL)* | **398** |
| *Total* | 400 |

3. Pause/stop the sequencing run (keep flow cell in device)
4. Ensure flow cell priming port and SpotON sample port covers are closed

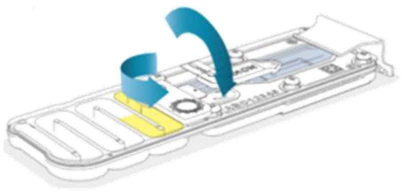

5. Remove waste buffers with P1000 pipet
   5.1. Set pipet to 1000 µL
   5.2. Insert in waste port 1
   5.3. Slowly aspirate waste buffer

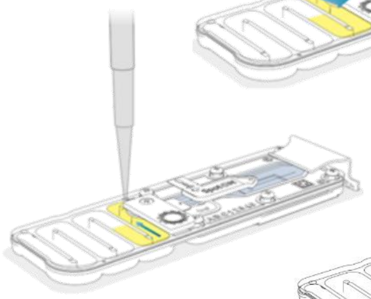

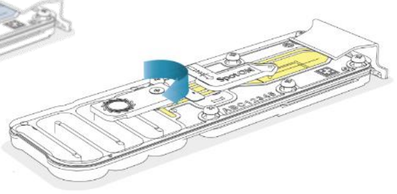

6. Slide flow cell priming port cover clockwise open
7. Set P1000 to 200µL, insert tip into priming port, turn **pipet volume up to max 220-230µL**, or until volume in tip

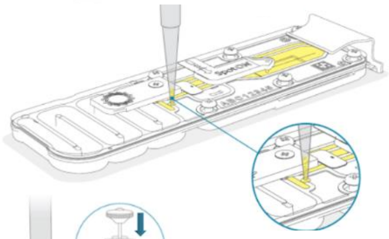

8. Load 200 µL of Flow Cell Wash Mix
   8.1. Set P1000 to **400 µL** take Flow Cell Wash Mix
   8.2. Insert tip into priming port, turn pipet **volume down to 200µL** by twisting pipet wheel slowly
   8.3. Slow but continuous rate!
   8.4. Incubate for 5 minutes and repeat with remaining 200 µL (push plunger slow and steady)
   8.5. Close priming port cover and incubate **1 hour**

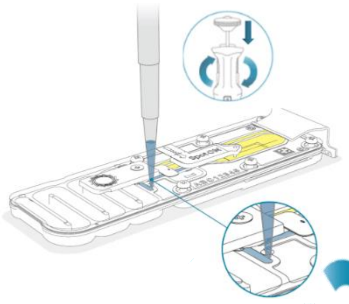

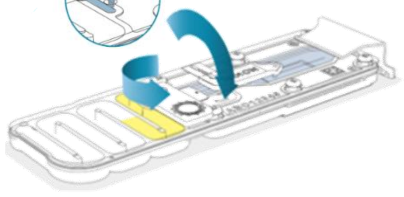

9. **After 1 hour!** Remove waste buffers with P1000
   9.1. Set pipet to 1000 µL, insert in waste port 1, slowly aspirate waste buffer
10. Flow cell is washed, **2 options**
    10.1. Run second sequencing library (**Priming and loading**)
    10.2. Store flow cell

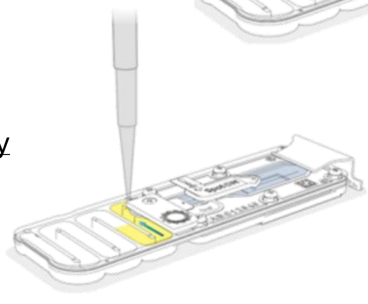

### I. Store flow cell

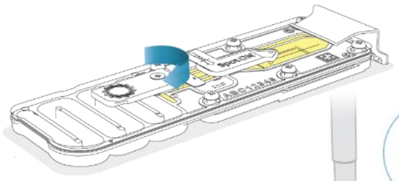

1. Thaw Storage Buffer (SB) at room temperature

2. Slide flow cell priming port cover clockwise open

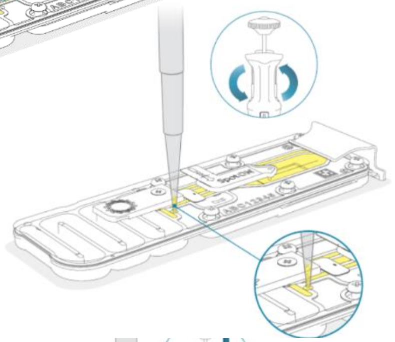

3. Set P1000 to 200µL, insert tip into priming port, turn **pipet volume up to max 220-230µL**, or until volume in tip

4. Load 500 µL of Storage Buffer (SB) into priming port (>5 sec)
   4.1. Set P1000 **to 700 µL** take Storage Buffer
   4.2. Insert tip into priming port, turn pipet volume **down to 200µL** by twisting pipet wheel in a slow but continuous rate!
   4.3. Close priming port cover

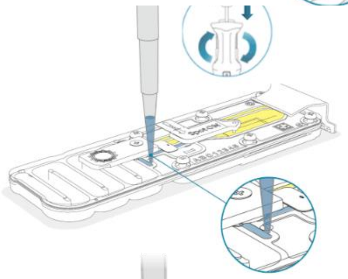

5. Remove waste buffers with P1000
   5.1. Set pipet to 1000 µL, insert in waste port 1, slowly aspirate waste buffer

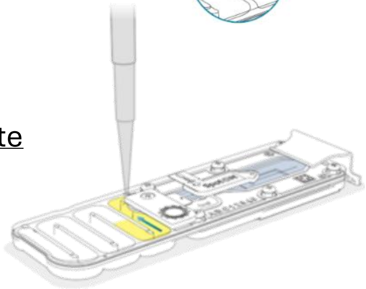

6. Flow cell can be stored at 2-8°C
