## Supplementary material for "Blood-meal analysis indicates preferential feeding on humans and non-commensal rodents by *Culex* mosquitoes in Tanzania": Bioinformatics: Bioinformatics.html

Supplementary Information: Bioinformatics


#### Table of contents

- 1 Analysis summary
  - 1.1 Basecalling and demultiplexing
  - 1.2 Putative chimeric reads
  - 1.3 Quality filtering
  - 1.4 Consensus sequence creation
  - 1.5 Taxonomic assignment
  - 1.6 Contamination correction and filtering
  - 1.7 References
- 2 Analysis scripts
  - 2.1 Script: Basecalling and demultiplexing
  - 2.2 Script: Putative chimeric reads
  - 2.3 Script: Python pipeline
  - 2.4 Script: Quality filtering
  - 2.5 Script: Consensus sequence creation
  - 2.6 Script: Taxonomic assignment

### Supplementary Information: Bioinformatics

Blood-meal analysis indicates preferential feeding on humans and non-commensal rodents by Culex mosquitoes in Tanzania

Author

Wim De Kesel

All analyses were conducted in reproducible, module-controlled environments on the University of Antwerp HPC infrastructure or a local workstation.

#### 1 Analysis summary

##### 1.1 Basecalling and demultiplexing

Oxford Nanopore POD5 files were processed on the University of Antwerp high-performance computing (HPC) cluster using SLURM. Basecalling was performed with Dorado v0.9.1 (Oxford Nanopore Technologies, 2023) using the super-accuracy model (dna\_r10.4.1\_e8.2\_400bps\_sup@v5.0.0) on GPU-enabled nodes. During basecalling, sequencing adapters were trimmed while barcode sequences were retained. Resulting BAM files were demultiplexed with Dorado using the SQK-NBD114-96 native barcoding kit, and sample-specific FASTQ files were generated. Barcode trimming was performed during the demultiplexing step.

##### 1.2 Putative chimeric reads

Demultiplexed FASTQ files were processed on the University of Antwerp HPC system using SLURM. A custom Python pipeline was executed in parallel across samples using GNU Parallel (32 concurrent jobs) to identify and split putative chimeric reads based on primer-binding sites

Chimeric reads were identified using a custom Python script implemented with Biopython and the regex package. The script searched each read for occurrences of the expected forward and reverse primer sequences, including reverse-complement orientations, allowing up to three mismatches per primer match. Reads containing internal primer-binding sites were split at the detected primer locations, and resulting sequence fragments were retained with their corresponding quality scores and written to new FASTQ files for downstream analysis.

##### 1.3 Quality filtering

Quality control filtering was performed using Chopper/NanoPack2 v0.9.0 (De Coster & Rademakers, 2023) on chimera-split FASTQ files. Reads with a minimum length of 70 bp and a mean Phred quality score of at least 15 were retained for downstream analyses. Reads failing either threshold were written to separate output files, while reads passing both criteria were retained in the final dataset. Empty output files generated during filtering were removed.

##### 1.4 Consensus sequence creation

Reads were first binned as likely 12S rRNA or COX1 amplicon sequences based on read length (12S: 150 ± 70 bp; COX1: 650 ± 70 bp). Within each amplicon class, reads were clustered based on sequence similarity and consensus sequences were generated using NGSpeciesID v0.3.0 (Sahlin et al., 2021), which utilizes SPOA v4.1.0 (Vaser et al., 2017) for sequence alignment and Medaka v1.11.3 (Oxford Nanopore Technologies, n.d.) for consensus polishing. Clustering sensitivity was normalized per sample by scaling the abundance ratio according to the number of reads within the expected amplicon length range, corresponding to a minimum cluster size of approximately 10 reads. For 12S sequences, stringent clustering parameters were applied (aligned\_threshold = 0.99, mapped\_threshold = 0.99, min\_fraction = 0.99), whereas for the more variable COX1 amplicons less stringent thresholds were used (aligned\_threshold = 0.90, mapped\_threshold = 0.90, min\_fraction = 0.90). More stringent clustering parameters were applied for the host identification to avoid clustering closely related species. Primer sequences were trimmed during consensus generation, and polished 12S and COX1 consensus sequences were combined per sample for downstream analyses.

##### 1.5 Taxonomic assignment

Consensus sequences were taxonomically assigned using a custom BLASTn-based (Camacho et al., 2009) pipeline against a locally curated mitochondrial reference database containing Culicidae and tetrapod sequences obtained from GenBank (downloaded 30 January 2025), supplemented with mitochondrial sequences from locally occurring species lacking suitable reference sequences. For each query sequence, up to ten database matches were retrieved and filtered using a minimum threshold of 90% sequence identity and 90% query coverage. Taxonomic assignment was based on the highest BLAST bit score. When multiple hits shared the highest bit score, a lowest common ancestor (LCA) was determined using the ETE3 (Huerta-Cepas et al., 2016) implementation of the NCBI taxonomy database. Results were compiled into a single CSV file containing taxonomic assignments and associated alignment statistics. Assignments based on less than 100% sequence identity or query coverage were manually verified against the GenBank core nucleotide (nt) database to assess potential species-level misidentifications and confirm marker-specific taxonomic assignments.

##### 1.6 Contamination correction and filtering

To account for background contamination introduced during DNA extraction, consensus sequences obtained from each mosquito specimen were compared with the negative control of the corresponding DNA extraction batch. Comparisons were based on both (i) the taxonomic identity assigned to each consensus sequence (host or vector) and (ii) the number of reads supporting that consensus sequence. For each host or vector taxon detected in a sample, the number of supporting reads observed in the corresponding negative extraction control was subtracted from the read support of matching consensus sequences in the sample. When a given taxon was not detected in the extraction negative control, a default correction value of 10 supporting reads was applied. This approach was used to reduce the influence of low-level background contamination introduced during DNA extraction.

Example 1:

Sample X:

- Taxon: Homo sapiens
- Consensus support: 250 reads
- Corresponding negative control: Homo sapiens - 30 reads
- Corrected consensus support: 220 reads (250-30)

Sample Y:

- Taxon: Corvus spp.
- Consensus support: 80 reads
- Corresponding negative control: Corvus spp. - not detected –> default 10 reads
- Corrected consensus support: 70 reads (70-10)

In example 1, 30 supporting reads assigned to Homo sapiens were detected in the extraction negative control and subtracted from sample X. For Corvus spp., the taxon was absent from the negative control, and the default correction value of 10 reads was applied.

To account for potential barcode hopping and cross-contamination during library preparation and sequencing, PCR negative controls were evaluated separately. For each taxonomic group (hosts and vectors independently), a detection threshold was calculated as the maximum number of supporting reads observed among the PCR negative controls of that respective sequencing run plus one standard deviation. To avoid excessively permissive thresholds, a minimum threshold of 100 supporting reads was applied irrespective of the value calculated from the negative controls. Consensus sequences with support below the resulting threshold were discarded from downstream analyses.

Exampe 2:

Suppose the highest support observed for any host taxon in the PCR negative controls was 65 reads, with a standard deviation of 20 reads:

```
Threshold ==> 65 + 20 = 85 reads
```

Because the pipeline imposes a minimum threshold of 100 reads, the effective threshold becomes:

```
max(85,100) = 100 reads
```

Any host consensus sequence supported by fewer than 100 reads would therefore be excluded from further analyses.

#### 2 Analysis scripts

##### 2.1 Script: Basecalling and demultiplexing

```
#!/bin/bash

#SBATCH --job-name=basecalling
#SBATCH --ntasks=1
#SBATCH --chdir= directory where files are stored
#SBATCH --partition=ampere_gpu
#SBATCH --gpus-per-node=2
#SBATCH --time=04:00:00
#SBATCH --account= account ID
#SBATCH --mail-type=ALL

#As an example: running this code with 2 gpus on approximately 11 million reads in super basecalling mode takes 02:30:00

module purge
module load calcua/2023a
module load dorado/0.9.1-foss-2023a-CUDA-12.1.1

#Make directory to store the basecalling and demultiplexing
mkdir -p ./basecalling/barcodes/demux

#Dorado basecaller (models are installed in private data account, you can download them from the Nanopore website), when specifying the pod5 folder it uses all the files within that folder, trim the adapters but not the barcodes
dorado basecaller /data/antwerpen/210/vsc21035/DoradoModels/dna_r10.4.1_e8.2_400bps_sup@v5.0.0 ./pod5_skip --trim adapters --output-dir ./basecalling

#Dorado demultiplexing, use the bam file create from code above, save in the created directories, save the output in fastq format and the used barcoding kit name
dorado demux ./basecalling/*.bam --output-dir ./basecalling/barcodes/demux --emit-fastq --kit-name SQK-NBD114-96
```

##### 2.2 Script: Putative chimeric reads

```
#!/bin/bash

#SBATCH --job-name=ChimeraSplit
#SBATCH --ntasks=1
#SBATCH --chdir= directory where files are stored
#SBATCH --cpus-per-task=64
#SBATCH --time=01:00:00
#SBATCH --account= account ID
#SBATCH --mail-type=ALL

# Load a Python module
module purge
module load calcua/2023a
module load calcua/all
module spider Python/3.11.3-GCCcore-12.3.0
module load parallel/20250722-GCCcore-14.2.0

# Directory containing FASTQ files
INPUT_DIR="./basecalling/barcodes/demux"

# Exclude unclassified file
FILES=$(ls $INPUT_DIR/*.fastq | grep -v "unclassified.fastq")

# Run parallel
parallel -j 32 "python ~/split_chimeras.py {}" ::: $FILES
```

##### 2.3 Script: Python pipeline

```
#!/usr/bin/env python3

import os
import sys
import regex
from Bio import SeqIO
from Bio.Seq import Seq

# --- SETTINGS ---
MAX_MISMATCHES = 3

# Primers
FORWARD_PRIMERS = {
    "LCO1490": "GGTCAACAAATCATAAAGATATTGG",
    "12S_V5f": "ACTGGGATTAGATACCCC"
}

REVERSE_PRIMERS = {
    "HCO2198": "TAAACTTCAGGGTGACCAAAAAATCA",
    "12S_V5r": "TAGAACAGGCTCCTCTAG"
}

def rc(seq):
    return str(Seq(seq).reverse_complement())

# Prepare orientations
PRIMERS = []
for name, seq in FORWARD_PRIMERS.items():
    PRIMERS.append((seq, 'forward', False, name))
    PRIMERS.append((rc(seq), 'forward', True, name))
for name, seq in REVERSE_PRIMERS.items():
    PRIMERS.append((seq, 'reverse', False, name))
    PRIMERS.append((rc(seq), 'reverse', True, name))

def find_all_primers(seq):
    hits = []
    for pseq, ptype, is_rc, name in PRIMERS:
        pattern = f"({pseq}){{s<={MAX_MISMATCHES}}}"
        for m in regex.finditer(pattern, seq, overlapped=True):
            start, end = m.span()
            hits.append((start, end, ptype, is_rc, name))
    hits.sort(key=lambda x: x[0])
    return hits

def split_by_primers(record):
    seq = str(record.seq)
    qual = record.letter_annotations["phred_quality"]
    hits = find_all_primers(seq)
    fragments = []
    last_idx = 0

    for start, end, ptype, is_rc, name in hits:
        if start < last_idx:
            continue
        cut_pos = start if not is_rc else end
        if cut_pos > last_idx:
            frag_seq = seq[last_idx:cut_pos]
            frag_qual = qual[last_idx:cut_pos]
            fragments.append(SeqIO.SeqRecord(
                Seq(frag_seq),
                id=f"{record.id}_frag_{last_idx}_{cut_pos}",
                description=f"fragment before primer {name}",
                letter_annotations={"phred_quality": frag_qual}
            ))
        last_idx = cut_pos

    if last_idx < len(seq):
        frag_seq = seq[last_idx:]
        frag_qual = qual[last_idx:]
        fragments.append(SeqIO.SeqRecord(
            Seq(frag_seq),
            id=f"{record.id}_frag_{last_idx}_end",
            description="tail",
            letter_annotations={"phred_quality": frag_qual}
        ))

    return fragments


# --- MAIN ---
if len(sys.argv) != 2:
    print("Usage: split_chimeras.py <input_fastq>")
    sys.exit(1)

input_path = sys.argv[1]
fname = os.path.basename(input_path)

# Output next to the input file
output_dir = os.path.join(os.path.dirname(input_path), "chimerasplit")
os.makedirs(output_dir, exist_ok=True)

output_path = os.path.join(output_dir, fname)

print(f"Processing: {fname}")

with open(output_path, "w") as out_handle:
    for rec in SeqIO.parse(input_path, "fastq"):
        fragments = split_by_primers(rec)
        SeqIO.write(fragments, out_handle, "fastq")

print(f" → Saved: {output_path}")
```

##### 2.4 Script: Quality filtering

```
#!/bin/bash

#SBATCH --job-name=QC
#SBATCH --ntasks=1
#SBATCH --chdir= directory where files are stored
#SBATCH --cpus-per-task=64
#SBATCH --time=00:10:00
#SBATCH --account= account ID
#SBATCH --mail-type=ALL

module purge
module load calcua/2023a
module load chopper/0.9.0-GCCcore-12.3.0
module load parallel/20230722-GCCcore-12.3.0

#Make directories for pass and failed reads
mkdir -p ./basecalling/barcodes/pass ./basecalling/barcodes/fail

#Run the chopper command and save the output with the new file name in the pass and fail directory
parallel -j 16 'chopper --quality 15 --minlength 70 --input {} > ./basecalling/barcodes/pass/$(echo {} | sed "s/.*_//")' ::: ./basecalling/barcodes/demux/chimerasplit/*.fastq

parallel -j 16 'chopper --inverse --quality 15 --minlength 70 --input {} > ./basecalling/barcodes/fail/$(echo {} | sed "s/.*_//")' ::: ./basecalling/barcodes/demux/chimerasplit/*.fastq

#Remove all the files that are empty
find ./basecalling/barcodes -type f -empty -delete
```

##### 2.5 Script: Consensus sequence creation

```
#!/bin/bash

#SBATCH --job-name=Consensus
#SBATCH --ntasks=1
#SBATCH --chdir= directory where files are stored
#SBATCH --cpus-per-task=16
#SBATCH --time=04:00:00
#SBATCH --account= account ID
#SBATCH --array=1-96      #Set it to the range of samples you have or select one specific sample number

module purge
module load calcua/2023a
module load NGSpeciesID/0.3.0-foss-2023a
module load spoa/4.1.0-GCC-12.3.0
module spider medaka/1.11.3-foss-2023a

DIR="./basecalling/barcodes/pass"
PRIMERFILE="/user/antwerpen/210/vsc21035/PrimersMQTZ.fasta"

FASTQFILE=$(ls $DIR/*.fastq | sed -n "${SLURM_ARRAY_TASK_ID}p")
FASTQBASENAME=$(basename "$FASTQFILE" .fastq)

mkdir -p Consensus/$FASTQBASENAME/{12S,COX1}
mkdir -p Consensus/AllConsensus/Queries

echo "Started $FASTQBASENAME"

### 12S ###
MEANamplicon_12S=150
SDamplicon_12S=70
COUNT_12S=$(awk -v mean="$MEANamplicon_12S" -v sd="$SDamplicon_12S" \
    'NR % 4 == 2 { if (length($0)>=mean-sd && length($0)<=mean+sd) c++ } END { print c }' \
    "$FASTQFILE")
FRACTION_12S=$(echo "scale=6; 10 / $COUNT_12S" | bc)
NGSpeciesID --ont --m $MEANamplicon_12S --s $SDamplicon_12S \
    --abundance_ratio $FRACTION_12S --consensus \
    --t $SLURM_CPUS_PER_TASK --medaka \
    --fastq "$FASTQFILE" \
    --aligned_threshold 0.99 \
    --mapped_threshold 0.99 \
    --min_fraction 0.99 \
    --min_shared 15 \
    --k 11 \
    --w 10 \
    --outfolder Consensus/$FASTQBASENAME/12S \
    --primer_file $PRIMERFILE

### COX1 ###
MEANamplicon_COX1=650
SDamplicon_COX1=70
COUNT_COX1=$(awk -v mean="$MEANamplicon_COX1" -v sd="$SDamplicon_COX1" \
    'NR % 4 == 2 { if (length($0)>=mean-sd && length($0)<=mean+sd) c++ } END { print c }' \
    "$FASTQFILE")
FRACTION_COX1=$(echo "scale=6; 10 / $COUNT_COX1" | bc)

NGSpeciesID --ont --m $MEANamplicon_COX1 --s $SDamplicon_COX1 \
    --abundance_ratio $FRACTION_COX1 --consensus \
    --t $SLURM_CPUS_PER_TASK --medaka \
    --fastq "$FASTQFILE" \
    --aligned_threshold 0.9 \
    --mapped_threshold 0.9 \
    --min_fraction 0.9 \
    --min_shared 80 \
    --k 15 \
    --w 20 \
    --outfolder Consensus/$FASTQBASENAME/COX1 \
    --primer_file $PRIMERFILE
# Combine
cat Consensus/$FASTQBASENAME/12S/medaka_cl_id_*/consensus.fasta \
    Consensus/$FASTQBASENAME/COX1/medaka_cl_id_*/consensus.fasta \
    > Consensus/AllConsensus/Queries/$FASTQBASENAME-Consensus.fasta

echo "Completed $FASTQBASENAME"
```

##### 2.6 Script: Taxonomic assignment

```
#!/usr/bin/env python3
# Run this in the AllConsensus location (python ../../../RunFiles/blast_pipeline.py)
# make sure the database is correct, if you add new fasta files do the following
  # update the fast file with new sequences, update the description with taxids and species
  # update the taxidmap file with the name of the fastas and taxid id
  # run then this code in terminal makeblastdb -in ./RunFiles/RefSeqDB.fasta -dbtype nucl -out RefSeqDataBase/RefSeqDB -parse_seqids -taxid_map taxid_map.txt

"""
BLAST folder pipeline:
- multiple FASTA files
- best hit by bit score
- LCA for ties
- min %identity and %query coverage
- Excel-ready CSV output
"""

import subprocess
import pandas as pd
from ete3 import NCBITaxa
from pathlib import Path

# ---------------- CONFIGURATION ---------------- #
query_folder = "Queries"              # folder with FASTA files
blast_db = "/mnt/c/Users/wimde/Desktop/NanoporeResults/RefSeqDataBase/RefSeqDB"
output_csv = "final_results.csv"

max_target_seqs = 10
min_pident = 90
min_qcov = 90
threshold_sum = min_pident + min_qcov

# Initialize NCBI taxonomy
ncbi = NCBITaxa()

# BLAST output columns
cols = [
    "query", "qlen", "sseqid", "slen", "pident",
    "length", "qcovs", "evalue", "bitscore",
    "taxid", "sscinames"
]

# ---------- LCA FUNCTION ---------- #
def compute_lca(taxids):
    lineages = [ncbi.get_lineage(t) for t in taxids]
    min_len = min(len(l) for l in lineages)
    lca = None
    for i in range(min_len):
        vals = {l[i] for l in lineages}
        if len(vals) == 1:
            lca = vals.pop()
        else:
            break
    return lca

# ---------- COLLECT FINAL RESULTS ---------- #
final_rows = []

# ---------- PROCESS EACH FASTA ---------- #
for fasta in sorted(Path(query_folder).glob("*.fa*")):
    fasta_name = fasta.name
    blast_out = f"{fasta.stem}_blast.tsv"

    print(f"Processing {fasta_name}")

    # ---- Run BLAST ---- #
    blast_cmd = [
        "blastn",
        "-query", str(fasta),
        "-db", blast_db,
        "-max_target_seqs", str(max_target_seqs),
        "-outfmt",
        "6 qseqid qlen sseqid slen pident length qcovs evalue bitscore staxids sscinames",
        "-out", blast_out
    ]
    subprocess.run(blast_cmd, check=True)

    # ---- Load BLAST results ---- #
    df = pd.read_csv(blast_out, sep="\t", names=cols)

    if df.empty:
        continue

    # ---- Apply average filter ---- #
    df = df[(df["pident"] + df["qcovs"] >= threshold_sum)]

    if df.empty:
        continue

    # ---- Keep best hits by highest bit score ---- #
    df["max_bitscore"] = df.groupby("query")["bitscore"].transform("max")
    df_best = df[df["bitscore"] == df["max_bitscore"]].drop(columns="max_bitscore")

    # ---- Process each query ---- #
    for query, group in df_best.groupby("query"):
        taxids = group["taxid"].astype(int).tolist()
        query_value = query.split("_")[-1]

        if len(taxids) == 1:
            row = group.iloc[0]
            taxonomy = row["sscinames"]
            best_match = row["sseqid"]
        else:
            ancestor = compute_lca(taxids)
            taxonomy = ncbi.get_taxid_translator([ancestor])[ancestor]
            best_match = ";".join(group["sseqid"])

        row = group.iloc[0]
        final_rows.append({
            "Fasta_file": fasta_name,
            "Query": query,
            "Query_value": query_value,
            "Query_length": row["qlen"],
            "Best_match": best_match,
            "Percent_identity": row["pident"],
            "Query_coverage": row["qcovs"],
            "Taxonomy": taxonomy
        })

# ---------- WRITE FINAL CSV ---------- #
final_df = pd.DataFrame(final_rows)
final_df.to_csv(output_csv, index=False)

print(f"\nb Final combined results written to: {output_csv}")
```
