## Supplementary material for "Blood-meal analysis indicates preferential feeding on humans and non-commensal rodents by *Culex* mosquitoes in Tanzania": Mosquito analysis: Mosquito_analysis.html

###### Wim De Kesel

- 1 Loading packages
- 2 Retrieve data
- 3 Data formatting
  - 3.1 Host proportions
  - 3.2 Adding metadata
  - 3.3 Name alterations
  - 3.4 Write data out to excel
- 4 Exploratory analysis
- 5 Network construction
  - 5.1 General network analysis
    - 5.1.1 Data filtering and genus-level
      splitting
    - 5.1.2 Host–host network
      construction
    - 5.1.3 Host-host matrix
      construction
  - 5.2 Culex network analysis
    - 5.2.1 Sitesplitting
    - 5.2.2 Season splitting
- 6 Human feedings
  - 6.1 Filtering to human
    feedings
  - 6.2 Overall host model
  - 6.3 Full interaction model
  - 6.4 Host site model
  - 6.5 Host Season model

### 1 Loading packages

```
# Load all the needed libraries
library(readxl)
library(skimr)
library(stringr)
library(treemap)
library(d3treeR)
library(ggplot2)
library(RColorBrewer)
library(htmlwidgets)
library(stringr)
library(lmerTest)
library(emmeans)
library(tidyr)
library(lme4)
library(DHARMa)
library(emmeans)
library(multcomp)
library(gridExtra)
library(dplyr)
library(tidyverse)
library(conflicted)
library(ggh4x)
library(writexl)
library(igraph)
library(ggraph)
library(oaqc)
library(glmmTMB)
library(ineq)
library(ggnewscale)
library(seriation)
library(reshape2)
library(multcompView)
library(DT)
conflicts_prefer(dplyr::select)
conflicts_prefer(dplyr::filter)

load("C:/Users/wimde/OneDrive - Universiteit Antwerpen/Data&Results/Mosquitoes/MQTanzania/.RData")
setwd("C:/Users/wimde/OneDrive - Universiteit Antwerpen/Data&Results/Mosquitoes/MQTanzania")
```

### 2 Retrieve data

```
# Read data
DataSes1 <- as.data.frame(read_excel("LabSession1/ResultsSes1.xlsx", 
    sheet = "CombinedResultsSes1",guess_max = 5000))
DataSes1 <- DataSes1 %>% select(1:22)

DataSes2 <- as.data.frame(read_excel("LabSession2/ResultsSes2.xlsx", 
    sheet = "CombinedResultsSes2"))
DataSes2 <- DataSes2%>% select(1:22)

DataSes3 <- as.data.frame(read_excel("LabSession3/ResultsSes3.xlsx", 
    sheet = "CombinedResultsSes3"))
DataSes3 <- DataSes3%>% select(1:22)

DataSes4 <- as.data.frame(read_excel("LabSession4/ResultsSes4.xlsx", 
    sheet = "CombinedResultsSes4"))
DataSes4 <- DataSes4%>% select(1:22)

# Field data
FieldData <- as.data.frame(read_excel("C:/Users/wimde/OneDrive - Universiteit Antwerpen/Data&Results/Data fieldworkTZ/FIELDLIST_WimPhD.xlsx",sheet = "MQ_Captures", range = "A2:AE8930"))
```

### 3 Data formatting

#### 3.1 Host proportions

```
# Function to calculate host proportions for each mosquito and identify the species
process_session <- function(data) {
  
  lookup_table <- data %>%
    filter(HostType == "Mosquito", Contamination == "No", is.na(IfNotTargetGene)) %>%
    group_by(MQ) %>%
    slice_max(ReCalculatedQuery_value, n = 1, with_ties = FALSE) %>%
    select(MQ, Final_Taxonomy) %>%
    ungroup()
  
  filtered <- data %>%
    filter(HostType == "Host", Contamination == "No", is.na(IfNotTargetGene)) %>%
    select(Session, ExtractionBatch, MQ,
           ReCalculatedQuery_value, Final_Taxonomy) %>%
    left_join(lookup_table, by = "MQ") %>%
    rename(
      Host   = Final_Taxonomy.x,
      Vector = Final_Taxonomy.y,
      Query_value = ReCalculatedQuery_value
    ) %>%
    mutate(Vector = coalesce(Vector, "Uncertain")) %>%
    group_by(MQ) %>%
    mutate(HostProportion = Query_value /
             sum(Query_value, na.rm = TRUE),
           TotalReads = sum(Query_value, na.rm = TRUE)) %>%
    ungroup()
  
  filtered
}

MQID <- function(data) {
  
  lookup_table <- data %>%
    filter(HostType == "Mosquito", !str_starts(MQ, "N")) %>%
    group_by(MQ) %>%
    slice_max(ReCalculatedQuery_value, n = 1, with_ties = FALSE) %>%
    select(Barcode, Session, ExtractionBatch, MQ, 
           Contamination, Final_Taxonomy) %>%
    ungroup()
  
return(lookup_table)
}

# Running function on the results of each Nanopore session
HostResults <- list(DataSes1, DataSes2, DataSes3, DataSes4) %>%
  lapply(process_session) %>%
  bind_rows()

MQResults <- list(DataSes1, DataSes2, DataSes3, DataSes4) %>%
  lapply(MQID) %>%
  bind_rows()


# Create Host genus category
HostResults <- HostResults %>%
  mutate(HostGenus = word(Host, 1))


# Identify MQs that appear more than once
duplicated_MQ <- MQResults %>%
  count(MQ) %>%
  filter(n > 1) %>%
  pull(MQ)

# For each duplicated MQ, find the row with the lowest TotalReads
MQ_session_to_remove <- HostResults %>%
  filter(MQ %in% duplicated_MQ) %>%
  group_by(MQ) %>%
  slice_min(order_by = TotalReads, n = 1, with_ties = FALSE) %>%
  ungroup() %>%
  select(MQ, Session) %>%
  rename(Session_to_remove = Session)

# Remove the double MQ's
HostResults <- HostResults %>%
  anti_join(MQ_session_to_remove, by = c("MQ", "Session" = "Session_to_remove"))
MQResults <- MQResults %>%
  anti_join(MQ_session_to_remove, by = c("MQ", "Session" = "Session_to_remove"))


# Only keep the results where the host proportion sum of host genus is at least X% based on the non-bloodfed control test
FilterHost <- HostResults %>%
  group_by(MQ, HostGenus) %>%
  mutate(HostPropSum = sum(HostProportion)) %>%
  ungroup() %>%
  filter(HostPropSum >=0.13)
```

#### 3.2 Adding metadata

```
# Adding year and location based on MQ id
HostResults <- HostResults %>%
  left_join(
    FieldData %>%
      mutate(ua_id_nr = as.character(ua_id_nr)) %>%
      select(ua_id_nr, year, Location),
    by = c("MQ" = "ua_id_nr")
  )

FilterHost <- FilterHost %>%
  left_join(
    FieldData %>%
      mutate(ua_id_nr = as.character(ua_id_nr)) %>%
      select(ua_id_nr, year, Location),
    by = c("MQ" = "ua_id_nr")
  )

MQResults <- MQResults %>%
  left_join(
    FieldData %>%
      mutate(ua_id_nr = as.character(ua_id_nr)) %>%
      select(ua_id_nr, year, Location),
    by = c("MQ" = "ua_id_nr")
  )
```

#### 3.3 Name alterations

```
# Overview of the data and make alterations to the names
FilterHost %>%
  select(Host) %>%
  lapply(unique)
```

```
## $Host
##  [1] "Homo sapiens"              "Mastomys natalensis"      
##  [3] "Lophuromys flavopunctatus" "Crocidura"                
##  [5] "Corvidae"                  "Gallus gallus"            
##  [7] "Felis catus"               "Threskiornis aethiopicus" 
##  [9] "Chlorocebus"               "Crocidura dsinezumi"      
## [11] "Centropus superciliosus"   "Zyzomys argurus"          
## [13] "Corvus albus"              "Megaloglossus woermanni"  
## [15] "Mus musculus"              "Praomys jacksoni"         
## [17] "Philentoma pyrhoptera"     "Canis"                    
## [19] "Canis lupus familiaris"    "Bos"                      
## [21] "Rhynchocyon"               "Sus scrofa"               
## [23] "Homininae"                 "Gerbillinae"              
## [25] "Acomys cahirinus"          "Meriones"                 
## [27] "Rattus"                    "Passeroidea"              
## [29] "Sylvisorex lunaris"        "Meleagris gallopavo"      
## [31] "Rattus norvegicus"         "Bubulcus ibis"            
## [33] "Fregata magnificens"       "Ovis"                     
## [35] "Colius striatus"
```

```
FilterHost %>%
  select(HostGenus) %>%
  lapply(unique)
```

```
## $HostGenus
##  [1] "Homo"          "Mastomys"      "Lophuromys"    "Crocidura"    
##  [5] "Corvidae"      "Gallus"        "Felis"         "Threskiornis" 
##  [9] "Chlorocebus"   "Centropus"     "Zyzomys"       "Corvus"       
## [13] "Megaloglossus" "Mus"           "Praomys"       "Philentoma"   
## [17] "Canis"         "Bos"           "Rhynchocyon"   "Sus"          
## [21] "Homininae"     "Gerbillinae"   "Acomys"        "Meriones"     
## [25] "Rattus"        "Passeroidea"   "Sylvisorex"    "Meleagris"    
## [29] "Bubulcus"      "Fregata"       "Ovis"          "Colius"
```

```
FilterHost %>%
  select(Location) %>%
  lapply(unique)
```

```
## $Location
## [1] "Fence"        "Pest"         "House"        "Apopo"        "Fence_Fence" 
## [6] "Fence_Shamba" "Resthouse"    "Dam"
```

```
FilterHost %>%
  select(Vector) %>%
  lapply(unique)
```

```
## $Vector
##  [1] "Culex pipiens pipiens"  "Aedes aegypti"          "Culex nebulosus"       
##  [4] "Uncertain"              "Culex hutchinsoni"      "Culex neavei"          
##  [7] "Aedes simpsoni"         "Anopheles rivulorum"    "Culex pipiens molestus"
## [10] "Culex bitaeniorhynchus" "Mansonia africana"
```

```
# Change the uncertain vectors to the identification of the field list
FilterHost <- FilterHost %>%
  mutate(across(c(MQ), as.numeric))

FilterHost <- FilterHost %>%
  left_join(FieldData %>% select(ua_id_nr, genus_field),
            by = c("MQ" = "ua_id_nr")) %>%
  mutate(Vector = ifelse(Vector == "Uncertain",
                         ifelse(genus_field == "Unknown", "Uncertain", genus_field),
                         Vector)) %>%
  select(-genus_field)


FilterHost <- FilterHost %>%
  mutate(
    Site = case_when(
      str_starts(Location, "Fence") ~ "Rural",
      Location == "House" ~ "Urban",
      Location == "Dam" ~ "Rural",
      Location %in% c("Apopo", "Resthouse", "Pest") ~ "Suburban",
      TRUE ~ NA_character_
    )
  )%>%
  mutate(
    VectorGenus = case_when(
      str_starts(Vector, "Culex") ~ "Culex",
      str_starts(Vector, "Aedes") ~ "Aedes",
      str_starts(Vector, "Anopheles") ~ "Anopheles",
      str_starts(Vector, "Mansonia") ~ "Mansonia",
      Vector == "Uncertain" ~ "Uncertain",
      TRUE ~ NA_character_
    )
  )%>%
  mutate(
    Season = case_when(
      year == "2023" ~ "Wet",
      year == "2025" ~ "Dry",
      TRUE ~ NA_character_
    )
  )

# Create host group categories
FilterHost <- FilterHost %>%
  mutate(
    HostGroup = case_when(
  # Humans
  Host %in% c("Homininae", "Homo sapiens") ~ "Human",
  
  # Domestic mammals
  Host %in% c("Felis catus", "Canis", "Canis lupus familiaris", "Bos", "Sus scrofa", "Ovis") ~ "Domestic mammal",
  
  # Domestic birds
  Host %in% c("Gallus gallus", "Meleagris gallopavo") ~ "Domestic bird",
  
  # Wild mammals: primates
  Host %in% c("Chlorocebus") ~ "Non human primate",
  
  # Wild mammals: commensal Rodents
  Host %in% c("Mus musculus", "Rattus", "Rattus norvegicus") ~ "Commensal rodent",
  
  # Wild mammals: Non commensal Rodent
  Host %in% c("Acomys cahirinus", "Mastomys natalensis", "Meriones", "Lophuromys flavopunctatus", "Zyzomys argurus", "Praomys jacksoni", "Gerbillinae") ~ "Non commensal rodent",
  
  # Wild mammals: shrews
  Host %in% c("Crocidura", "Crocidura dsinezumi", "Sylvisorex lunaris") ~ "Shrew",
  
  # Wild mammals: other
  Host %in% c("Megaloglossus woermanni","Rhynchocyon") ~ "Wild mammal",
  
  # Wild birds: corvids
  Host %in% c("Corvidae","Corvus albus") ~ "Crow",
  
  # Wild birds: waterbirds / waders
  Host %in% c("Bubulcus ibis","Threskiornis aethiopicus","Fregata magnificens") ~
    "Water bird",
  
  # Wild birds: song birds
  Host %in% c("Philentoma pyrhoptera","Passeroidea","Colius striatus","Centropus superciliosus") ~
    "Other bird",
  TRUE ~ "Unknown"
))    

# Create broad host categories
FilterHost <- FilterHost %>%
  mutate(
    HostBroad = case_when(
  # Primates
  HostGroup %in% c("Human", "Non human primate") ~ "Primate",
  
  # Domestic animal
  HostGroup %in% c("Domestic mammal", "Domestic bird") ~ "Domestic animal",
  
  # Birds
  HostGroup %in% c("Crow", "Water bird", "Other bird") ~ "Bird",
  
  # Small mammal
  HostGroup %in% c("Commensal rodent", "Non commensal rodent", "Shrew", "Wild mammal") ~ "Small mammal",

  TRUE ~ "Unknown"
))  

# MQ results
# Change the uncertain vectors to the identification of the field list
MQResults <- MQResults %>%
  mutate(across(c(MQ), as.numeric))

MQResults <- MQResults %>%
  left_join(FieldData %>% select(ua_id_nr, genus_field),
            by = c("MQ" = "ua_id_nr")) %>%
  mutate(Final_Taxonomy = ifelse(Contamination == "Yes",
                         ifelse(genus_field == "Unknown", "Uncertain", genus_field),
                         Final_Taxonomy)) %>%
  select(-genus_field)

# Overview of the data and make alterations to the names
MQResults %>%
  select(Final_Taxonomy) %>%
  lapply(unique)
```

```
## $Final_Taxonomy
##  [1] "Aedes aegypti"          "Culex pipiens pipiens"  "Culex nebulosus"       
##  [4] "Culex neavei"           "Culex"                  "Culex hutchinsoni"     
##  [7] "Aedes simpsoni"         "Anopheles rivulorum"    "Mansonia"              
## [10] "Culex cinereus"         "Culex pipiens molestus" "Culex bitaeniorhynchus"
## [13] "Mansonia africana"
```

```
MQResults <- MQResults %>%
  mutate(
    Site = case_when(
      str_starts(Location, "Fence") ~ "Rural",
      Location == "House" ~ "Urban",
      Location == "Dam" ~ "Rural",
      Location %in% c("Apopo", "Resthouse", "Pest") ~ "Suburban",
      TRUE ~ NA_character_
    )
  )%>%
  mutate(
    VectorGenus = case_when(
      str_starts(Final_Taxonomy, "Culex") ~ "Culex",
      str_starts(Final_Taxonomy, "Aedes") ~ "Aedes",
      str_starts(Final_Taxonomy, "Anopheles") ~ "Anopheles",
      str_starts(Final_Taxonomy, "Mansonia") ~ "Mansonia",
      Final_Taxonomy == "Uncertain" ~ "Uncertain",
      TRUE ~ NA_character_
    )
  )%>%
  mutate(
    Season = case_when(
      year == "2023" ~ "Wet",
      year == "2025" ~ "Dry",
      TRUE ~ NA_character_
    )
  )
```

#### 3.4 Write data out to excel

```
# Write the data frames to excel files
write_xlsx(HostResults, "HostResults.xlsx") 

write_xlsx(FilterHost, "FilterHost.xlsx")    

write_xlsx(MQResults, "MQResults.xlsx")
```

### 4 Exploratory analysis

```
# Field data
FieldData <- FieldData %>%
  mutate(
    Site = case_when(
      str_starts(Location, "Fence") ~ "Rural",
      Location == "House" ~ "Urban",
      Location == "Dam" ~ "Rural",
      Location %in% c("Apopo", "Resthouse", "Pest") ~ "Suburban",
      TRUE ~ NA_character_
    )
  )%>%
  mutate(
    Season = case_when(
      year == "2023" ~ "Wet",
      year == "2025" ~ "Dry",
      TRUE ~ NA_character_
    )
  )

FieldData <- FieldData %>%
  filter(!is.na(Site))

# Data summary field data
FieldData <- FieldData %>%
  mutate(across(c(sex, bloodfed, genus_field, species_field, year, month, day, Season, Site), as.factor))

skim(FieldData)
```

Data summary

|  |  |
| --- | --- |
| Name | FieldData |
| Number of rows | 8917 |
| Number of columns | 32 |
| \_\_\_\_\_\_\_\_\_\_\_\_\_\_\_\_\_\_\_\_\_\_\_ |  |
| Column type frequency: |  |
| character | 18 |
| factor | 9 |
| logical | 3 |
| numeric | 2 |
| \_\_\_\_\_\_\_\_\_\_\_\_\_\_\_\_\_\_\_\_\_\_\_\_ |  |
| Group variables | None |

**Variable type: character**

| skim\_variable | n\_missing | complete\_rate | min | max | empty | n\_unique | whitespace |
| --- | --- | --- | --- | --- | --- | --- | --- |
| project\_code | 0 | 1.00 | 3 | 3 | 0 | 1 | 0 |
| ua\_id\_pr | 0 | 1.00 | 2 | 2 | 0 | 1 | 0 |
| ua\_id | 0 | 1.00 | 3 | 7 | 0 | 8917 | 0 |
| location\_code | 0 | 1.00 | 2 | 3 | 0 | 10 | 0 |
| session\_code | 3019 | 0.66 | 2 | 2 | 0 | 5 | 0 |
| Location | 0 | 1.00 | 3 | 12 | 0 | 9 | 0 |
| locses | 1 | 1.00 | 2 | 4 | 0 | 30 | 0 |
| Session\_check | 3020 | 0.66 | 4 | 4 | 0 | 21 | 0 |
| Classis | 0 | 1.00 | 7 | 7 | 0 | 1 | 0 |
| trap | 0 | 1.00 | 1 | 1 | 0 | 1 | 0 |
| trap\_id | 0 | 1.00 | 1 | 10 | 0 | 18 | 0 |
| Specimen intact? | 0 | 1.00 | 2 | 3 | 0 | 2 | 0 |
| IMG | 8865 | 0.01 | 5 | 26 | 0 | 51 | 0 |
| capture\_remarks | 8662 | 0.03 | 4 | 55 | 0 | 103 | 0 |
| epp\_nr | 5012 | 0.44 | 4 | 8 | 0 | 419 | 0 |
| plate\_ID | 3905 | 0.56 | 6 | 7 | 0 | 54 | 0 |
| plate\_row | 3905 | 0.56 | 1 | 1 | 0 | 8 | 0 |
| Position\_plate | 3905 | 0.56 | 8 | 10 | 0 | 5012 | 0 |

**Variable type: factor**

| skim\_variable | n\_missing | complete\_rate | ordered | n\_unique | top\_counts |
| --- | --- | --- | --- | --- | --- |
| day | 0 | 1 | FALSE | 24 | 1: 1124, 23: 787, 30: 684, 11: 559 |
| month | 0 | 1 | FALSE | 4 | 5: 5261, 9: 2338, 8: 681, 6: 637 |
| year | 0 | 1 | FALSE | 2 | 202: 5898, 202: 3019 |
| genus\_field | 0 | 1 | FALSE | 6 | Cul: 8325, Aed: 390, Man: 141, Unk: 46 |
| species\_field | 0 | 1 | FALSE | 25 | pip: 6734, sp: 1015, per: 683, aeg: 150 |
| sex | 0 | 1 | FALSE | 3 | F: 4949, M: 3907, Unk: 61 |
| bloodfed | 0 | 1 | FALSE | 3 | No: 8554, Yes: 227, Egg: 136 |
| Site | 0 | 1 | FALSE | 3 | Urb: 6201, Rur: 1690, Sub: 1026 |
| Season | 0 | 1 | FALSE | 2 | Wet: 5898, Dry: 3019 |

**Variable type: logical**

| skim\_variable | n\_missing | complete\_rate | mean | count |
| --- | --- | --- | --- | --- |
| id\_ext | 8917 | 0.00 | NaN | : |
| Day/Night | 8917 | 0.00 | NaN | : |
| HardCopy\_page | 5898 | 0.34 | 1 | TRU: 3019 |

**Variable type: numeric**

| skim\_variable | n\_missing | complete\_rate | mean | sd | p0 | p25 | p50 | p75 | p100 | hist |
| --- | --- | --- | --- | --- | --- | --- | --- | --- | --- | --- |
| ua\_id\_nr | 75 | 0.99 | 4431.82 | 2563.38 | 1 | 2212.25 | 4423.5 | 6661.75 | 8873 | ▇▇▇▇▇ |
| plate\_column | 3905 | 0.56 | 6.43 | 3.45 | 1 | 3.00 | 6.0 | 9.00 | 12 | ▇▅▅▅▇ |

```
Field_summary <- FieldData %>%
  group_by(genus_field, Site, Season, sex) %>%
  summarise(count = n_distinct(ua_id), .groups = "drop") %>%
  mutate(Season_Sex = interaction(Season, sex, sep = " & "))


Field_summary$Season_Sex <- factor(
  Field_summary$Season_Sex,
  levels = c("Dry & F", "Dry & M", "Dry & Unknown",
             "Wet & F", "Wet & M", "Wet & Unknown")
)

season_colors <- c(
  "Dry & F" = "#d95f02",
  "Dry & M"   = "#fddbc7",
  "Dry & Unknown" = "grey",
  "Wet & F" = "#7570b3",
  "Wet & M"   = "#dadaeb",
  "Wet & Unknown" = "grey"
)

CapturePlot <- ggplot(Field_summary,
       aes(x = Site, y = count, fill = Season_Sex, group = Season)) + 
  geom_bar(stat = "identity", position = position_dodge2(preserve = "single")) +
  facet_wrap(~ genus_field, scales = "free_y") +
  scale_fill_manual(values = season_colors,
                    name = "Season & Sex") +
  theme_minimal(base_size = 20) +
theme(strip.text = element_text(face = "italic")) +
  labs(
    title = "Total Captured Mosquitoes by Site and Season",
    y = "Count"
  )

ggsave(
    filename = "MQCapturePlot.svg",
    plot = CapturePlot,
    width = 14,
    height = 8, dpi = 600)
```

```
## Table of captured mosquito genera per location, season and sex
```

```
# Data summary MQ results
MQResults <- MQResults %>%
  mutate(across(c(MQ, Site, ExtractionBatch, year, Season, Session, VectorGenus), as.factor))
skim(MQResults)
```

Data summary

|  |  |
| --- | --- |
| Name | MQResults |
| Number of rows | 331 |
| Number of columns | 11 |
| \_\_\_\_\_\_\_\_\_\_\_\_\_\_\_\_\_\_\_\_\_\_\_ |  |
| Column type frequency: |  |
| character | 4 |
| factor | 7 |
| \_\_\_\_\_\_\_\_\_\_\_\_\_\_\_\_\_\_\_\_\_\_\_\_ |  |
| Group variables | None |

**Variable type: character**

| skim\_variable | n\_missing | complete\_rate | min | max | empty | n\_unique | whitespace |
| --- | --- | --- | --- | --- | --- | --- | --- |
| Barcode | 0 | 1 | 9 | 9 | 0 | 95 | 0 |
| Contamination | 0 | 1 | 2 | 3 | 0 | 2 | 0 |
| Final\_Taxonomy | 0 | 1 | 5 | 22 | 0 | 13 | 0 |
| Location | 0 | 1 | 3 | 12 | 0 | 8 | 0 |

**Variable type: factor**

| skim\_variable | n\_missing | complete\_rate | ordered | n\_unique | top\_counts |
| --- | --- | --- | --- | --- | --- |
| Session | 0 | 1 | FALSE | 4 | Ses: 90, Ses: 89, Ses: 79, Ses: 73 |
| ExtractionBatch | 0 | 1 | FALSE | 16 | Bat: 23, Bat: 23, Bat: 23, Bat: 23 |
| MQ | 0 | 1 | FALSE | 331 | 79: 1, 128: 1, 214: 1, 217: 1 |
| year | 0 | 1 | FALSE | 2 | 202: 245, 202: 86 |
| Site | 0 | 1 | FALSE | 3 | Urb: 273, Sub: 34, Rur: 24 |
| VectorGenus | 0 | 1 | FALSE | 4 | Cul: 322, Aed: 5, Man: 3, Ano: 1 |
| Season | 0 | 1 | FALSE | 2 | Wet: 245, Dry: 86 |

```
  # 331 unique MQ
  # 13 unique vectors (Final Taxonomy) - 4 genera

# Show how many mosquitoes for each distinct mosquito species were sequenced
MQSpeciesSeq <- MQResults %>%
  distinct(Final_Taxonomy, MQ) %>%     # keep only unique MQ per Vector
  count(Final_Taxonomy)

# Show how many mosquitoes for each distinct mosquito genus were sequenced
MQGenusQSeq <- MQResults %>%
  distinct(VectorGenus, MQ) %>%     # keep only unique MQ per Vector
  count(VectorGenus)


# Data summary blood results
FilterHost <- FilterHost %>%
  mutate(across(c(Site, ExtractionBatch, year, Season, Session, MQ), as.factor))

skim(FilterHost)
```

Data summary

|  |  |
| --- | --- |
| Name | FilterHost |
| Number of rows | 522 |
| Number of columns | 17 |
| \_\_\_\_\_\_\_\_\_\_\_\_\_\_\_\_\_\_\_\_\_\_\_ |  |
| Column type frequency: |  |
| character | 7 |
| factor | 6 |
| numeric | 4 |
| \_\_\_\_\_\_\_\_\_\_\_\_\_\_\_\_\_\_\_\_\_\_\_\_ |  |
| Group variables | None |

**Variable type: character**

| skim\_variable | n\_missing | complete\_rate | min | max | empty | n\_unique | whitespace |
| --- | --- | --- | --- | --- | --- | --- | --- |
| Host | 0 | 1 | 3 | 25 | 0 | 35 | 0 |
| Vector | 0 | 1 | 5 | 22 | 0 | 12 | 0 |
| HostGenus | 0 | 1 | 3 | 13 | 0 | 32 | 0 |
| Location | 0 | 1 | 3 | 12 | 0 | 8 | 0 |
| VectorGenus | 0 | 1 | 5 | 9 | 0 | 4 | 0 |
| HostGroup | 0 | 1 | 4 | 20 | 0 | 11 | 0 |
| HostBroad | 0 | 1 | 4 | 15 | 0 | 4 | 0 |

**Variable type: factor**

| skim\_variable | n\_missing | complete\_rate | ordered | n\_unique | top\_counts |
| --- | --- | --- | --- | --- | --- |
| Session | 0 | 1 | FALSE | 4 | Ses: 144, Ses: 132, Ses: 126, Ses: 120 |
| ExtractionBatch | 0 | 1 | FALSE | 16 | Bat: 51, Bat: 45, Bat: 42, Bat: 37 |
| MQ | 0 | 1 | FALSE | 310 | 597: 5, 783: 5, 886: 5, 278: 4 |
| year | 0 | 1 | FALSE | 2 | 202: 374, 202: 148 |
| Site | 0 | 1 | FALSE | 3 | Urb: 440, Sub: 52, Rur: 30 |
| Season | 0 | 1 | FALSE | 2 | Wet: 374, Dry: 148 |

**Variable type: numeric**

| skim\_variable | n\_missing | complete\_rate | mean | sd | p0 | p25 | p50 | p75 | p100 | hist |
| --- | --- | --- | --- | --- | --- | --- | --- | --- | --- | --- |
| Query\_value | 0 | 1 | 10489.46 | 22775.74 | 102.00 | 350.50 | 1144.50 | 7845.50 | 190126 | ▇▁▁▁▁ |
| HostProportion | 0 | 1 | 0.58 | 0.39 | 0.00 | 0.18 | 0.66 | 0.98 | 1 | ▅▂▂▂▇ |
| TotalReads | 0 | 1 | 24913.59 | 36078.86 | 118.00 | 1693.00 | 6575.00 | 39394.75 | 194537 | ▇▂▁▁▁ |
| HostPropSum | 0 | 1 | 0.76 | 0.30 | 0.13 | 0.55 | 0.96 | 1.00 | 1 | ▂▁▁▁▇ |

```
  # 310 unique MQ
  # 12 unique vectors - 4 genera
  # 35 unique hosts, circa 32 genera, 11 host groups, 4 broad groups
  # Reads/host: 10489 +- 22776, min=350, max=190126, median= 1144
  # HostReads/MQ: 24914 +- 36079, min=118, max=194537, median= 6575
  # HostProp/MQ: 0.578 +- 0.393, min=0.001, max=1.00, median=0.664
  # HostPropSum/MQ: 0.764 +- 0.302, min=0.131, max=1.00, median=0.958

# How many samples of each mosquito species with a blood meal result
MQSpeciesBlood <- FilterHost %>%
  distinct(Vector, MQ) %>%     # keep only unique MQ per Vector
  count(Vector)

# How many samples of each mosquito genus with a blood meal result
MQGenusBlood <- FilterHost %>%
  distinct(VectorGenus, MQ) %>%     # keep only unique MQ per Vector
  count(VectorGenus)

# Average number of used blood hosts per mosquito
HostsPerMQ <- FilterHost %>%
  group_by(MQ) %>%
  summarise(n_hosts = n_distinct(HostGroup)) %>%
  summarise(
    mean = round(mean(n_hosts),2),
    sd   = round(sd(n_hosts),2),
    min  = min(n_hosts),
    max  = max(n_hosts)
  )

# Average number of used blood hosts per mosquito genus
HostsPerGenus <- FilterHost %>%
  group_by(VectorGenus, MQ) %>%
  summarise(n_hosts = n_distinct(HostGroup), .groups = "drop") %>%
  group_by(VectorGenus) %>%
  summarise(mean = round(mean(n_hosts),2),
    sd   = round(sd(n_hosts),2),
    min  = min(n_hosts),
    max  = max(n_hosts)
  )
```

```
## Number of mosquitoes species that were sequenced
```

| Final\_Taxonomy | n |
| --- | --- |
| Aedes aegypti | 4 |
| Aedes simpsoni | 1 |
| Anopheles rivulorum | 1 |
| Culex | 15 |
| Culex bitaeniorhynchus | 1 |
| Culex cinereus | 1 |
| Culex hutchinsoni | 1 |
| Culex neavei | 4 |
| Culex nebulosus | 4 |
| Culex pipiens molestus | 1 |
| Culex pipiens pipiens | 295 |
| Mansonia | 1 |
| Mansonia africana | 2 |

```
## Number of mosquitoes per genus that were sequenced
```

| VectorGenus | n |
| --- | --- |
| Aedes | 5 |
| Anopheles | 1 |
| Culex | 322 |
| Mansonia | 3 |

```
## Number of mosquitoes species that had a blood meal result
```

| Vector | n |
| --- | --- |
| Aedes aegypti | 4 |
| Aedes simpsoni | 1 |
| Anopheles rivulorum | 1 |
| Culex | 11 |
| Culex bitaeniorhynchus | 1 |
| Culex hutchinsoni | 1 |
| Culex neavei | 3 |
| Culex nebulosus | 4 |
| Culex pipiens molestus | 1 |
| Culex pipiens pipiens | 280 |
| Mansonia | 1 |
| Mansonia africana | 2 |

```
## Number of mosquitoes per genus that had a blood meal result
```

| VectorGenus | n |
| --- | --- |
| Aedes | 5 |
| Anopheles | 1 |
| Culex | 301 |
| Mansonia | 3 |

```
## Mean number of detected host groups within all mosquitoes
```

| mean | sd | min | max |
| --- | --- | --- | --- |
| 1.35 | 0.62 | 1 | 4 |

```
## Mean number of detected host groups per mosquito genus
```

| VectorGenus | mean | sd | min | max |
| --- | --- | --- | --- | --- |
| Aedes | 1.20 | 0.45 | 1 | 2 |
| Anopheles | 1.00 | NA | 1 | 1 |
| Culex | 1.35 | 0.61 | 1 | 4 |
| Mansonia | 2.00 | 1.00 | 1 | 3 |

```
# Treemap of the used bloodhosts
p <- treemap(FilterHost, 
             index = c("HostBroad","HostGroup", "HostGenus","Host"),
             vSize = "HostProportion",
             type = "index",
             palette = "Set2",
             bg.labels=c("white"),
            align.labels=list(
              c("center", "center"), 
              c("right", "bottom")
            ))
```

```
(inter <- d3tree2( p ,  rootname = "Bloodhosts" ))
```

```
saveWidget(inter, file=paste0( getwd(), "/interactiveTreemap.html"))

# Proportion blood host per site and genus with sample size 
labels_site <- FilterHost %>%
  group_by(Site, VectorGenus) %>%
  summarise(N = n_distinct(MQ), .groups = "drop") %>%
  mutate(label = paste0(Site, "\n(N=", N, ")"))

scales_list <- lapply(
  split(labels_site, labels_site$VectorGenus),
  \(df) scale_x_discrete(breaks = df$Site, labels = df$label)
)

plot1 <- ggplot(FilterHost, aes(x = Site,
                        y = HostProportion,
                        fill = HostGroup)) +
  geom_bar(stat = "identity", position = "fill") +
  ggh4x::facet_wrap2(~VectorGenus, scales = "free_x") +
  ggh4x::facetted_pos_scales(x = scales_list) +
  scale_fill_manual(values = brewer.pal(12, "Set3")) +
  labs(y = "Host group proportion", fill = "Host group")
```

```
## Proportion of host groups per site and mosquito genus with sample size
```

```
# Proportion blood host per Season and genus with sample size 
labels_Season<- FilterHost %>%
  group_by(Season, VectorGenus) %>%
  summarise(N = n_distinct(MQ), .groups = "drop") %>%
  mutate(label = paste0(Season, "\n(N=", N, ")"))

scales_list <- lapply(
  split(labels_Season, labels_Season$VectorGenus),
  \(df) scale_x_discrete(breaks = df$Season, labels = df$label)
)

plot2 <- ggplot(FilterHost, aes(x = Season,
                        y = HostProportion,
                        fill = HostGroup)) +
  geom_bar(stat = "identity", position = "fill") +
  ggh4x::facet_wrap2(~VectorGenus, scales = "free_x") +
  ggh4x::facetted_pos_scales(x = scales_list) +
  scale_fill_manual(values = brewer.pal(12, "Set3")) +
  labs(y = "Host group proportion", fill = "Host group")
```

```
## Proportion of host group per season and mosquito genus with sample size
```

### 5 Network construction

#### 5.1 General network analysis

##### 5.1.1 Data filtering and genus-level splitting

```
# Filter to retain meaningful variables
NetworkData <- FilterHost %>%
  group_by(MQ, Site, HostGroup, HostBroad, Season, VectorGenus) %>%
  summarise(
    HostProportion = sum(HostProportion),
    .groups = "drop"
  )

# Split data by mosquito genus
genus_list <- split(NetworkData, NetworkData$VectorGenus)

# Count the number of mosquitoes for each genus
MQ_counts <- c(
  Aedes     = length(unique(genus_list$Aedes$MQ)),
  Anopheles = length(unique(genus_list$Anopheles$MQ)),
  Culex     = length(unique(genus_list$Culex$MQ)),
  Mansonia  = length(unique(genus_list$Mansonia$MQ))
)
```

##### 5.1.2 Host–host network construction

```
# Create empty list to store plots
network_plots <- list()

# Build SRI network
build_host_network_SRI <- function(df) {

  # Binary MQ × Host matrix
  mat <- df %>%
    mutate(presence = ifelse(HostProportion > 0, 1, 0)) %>%
    group_by(MQ, HostGroup) %>%
    summarise(value = max(presence), .groups = "drop") %>%
    pivot_wider(
      names_from = HostGroup,
      values_from = value,
      values_fill = 0
    )

  mat <- as.matrix(mat[, -1])
  rownames(mat) <- unique(df$MQ)

  # Number of MQ per host
  Xi <- colSums(mat)

  # Shared MQ between hosts
  Xij <- t(mat) %*% mat

  # Compute SRI matrix
  SRI <- matrix(0, nrow = ncol(mat), ncol = ncol(mat))
  colnames(SRI) <- colnames(mat)
  rownames(SRI) <- colnames(mat)

  for (i in 1:ncol(mat)) {
    for (j in 1:ncol(mat)) {
      if (i != j) {
        SRI[i, j] <- Xij[i, j] / (Xi[i] + Xi[j] - Xij[i, j])
      }
    }
  }

  diag(SRI) <- 0

  # Return BOTH graph and matrix
  list(
    graph = graph_from_adjacency_matrix(
      SRI,
      mode = "undirected",
      weighted = TRUE,
      diag = FALSE
    ),
    matrix = SRI
  )
}

# Create networks
host_networks <- lapply(genus_list, build_host_network_SRI)

# Store genus names
genus_names <- names(host_networks)

#Compute node proportions
host_mosquito_prop <- function(df) {
  df %>%
    distinct(MQ, HostGroup) %>%
    count(HostGroup, name = "N_MQ") %>%
    mutate(prop = N_MQ / length(unique(df$MQ)))
}

# Create network plots for each genus
for (genus in genus_names) {

  res <- host_networks[[genus]]
  g <- res$graph
  mat <- res$matrix   # <-- matrix available
  
  props <- host_mosquito_prop(genus_list[[genus]])

  V(g)$prop <- props$prop[match(V(g)$name, props$HostGroup)]
  V(g)$prop[is.na(V(g)$prop)] <- 0

  # Edge weights already SRI
  E(g)$weight <- as.numeric(E(g)$weight)

  # Plot
  p <- ggraph(g, layout = "fr") +
    geom_edge_link(
      aes(edge_color = weight),
      alpha = 0.8,
      width = 2.5
    ) +
    geom_node_point(
      aes(size = prop),
      colour = "#C11007"
    ) +
    geom_node_text(
      aes(label = name),
      repel = TRUE
    ) +
    scale_edge_color_gradient(low = "white",
      high = "#075BC1",limits = c(0, max(E(g)$weight))) +
                              
    labs(
      title = paste0("Feeding network of ", genus,
                     " (N= ", MQ_counts[genus], ")"),
      edge_color = "SRI",
      size = "Host group proportion"
    ) +
    theme_void()

  network_plots[[genus]] <- p
  print(p)
  
  ggsave(
    filename = paste0("FeedingNetwork_", genus, ".svg"),
    plot = p,
    width = 10,
    height = 6)
}
```

```
## Nodes represent host groups, with size indicating the proportion of mosquitoes feeding on each host group. Edges connect host groups that are fed upon by the same mosquitoes, with edge color reflecting the strength of this association calculated by Simple Ratio Index (SRI). It is a measure for how often two host groups are used together, relative to how often they are used overall.
```

##### 5.1.3 Host-host matrix construction

```
# Create a function to extract the matrix
build_host_matrix_counts <- function(df) {

  mat <- df %>%
    mutate(presence = ifelse(HostProportion > 0, 1, 0)) %>%
    group_by(MQ, HostGroup) %>%
    summarise(value = max(presence), .groups = "drop") %>%
    pivot_wider(
      names_from = HostGroup,
      values_from = value,
      values_fill = 0
    )

  mat <- as.matrix(mat[, -1])
  rownames(mat) <- unique(df$MQ)

  # Co-occurrence matrix
  Xij <- t(mat) %*% mat

  return(Xij)
}

build_host_matrices <- function(df) {

  mat <- df %>%
    mutate(presence = ifelse(HostProportion > 0, 1, 0)) %>%
    group_by(MQ, HostGroup) %>%
    summarise(value = max(presence), .groups = "drop") %>%
    pivot_wider(
      names_from = HostGroup,
      values_from = value,
      values_fill = 0
    )

  mat <- as.matrix(mat[, -1])
  rownames(mat) <- unique(df$MQ)

  # Counts
  Xij <- t(mat) %*% mat
  Xi <- diag(Xij)

  # SRI
  SRI <- matrix(0, ncol(mat), ncol(mat))
  colnames(SRI) <- colnames(mat)
  rownames(SRI) <- colnames(mat)

  for (i in 1:ncol(mat)) {
    for (j in 1:ncol(mat)) {
      if (i != j) {
        SRI[i, j] <- Xij[i, j] / (Xi[i] + Xi[j] - Xij[i, j])
      }
    }
  }

  # mcnemar
  MCnemarP <- matrix(1, ncol(mat), ncol(mat))
  colnames(MCnemarP) <- colnames(mat)
  rownames(MCnemarP) <- colnames(mat)
  
  for (i in 1:ncol(mat)) {
    for (j in 1:ncol(mat)) {
      if (i != j) {
        tab <- matrix(c(Xij[i, j], Xi[i]-Xij[i, j],Xi[j]-Xij[i, j],  301-Xi[i]-Xi[j]+Xij[i, j]), 
                      nrow = 2, byrow = TRUE)
        MCnemarP[i, j] <- round(p.adjust(mcnemar.test(tab)$p.value, method = "holm"),3)
      }
    }
  }
  MCnemarC <- matrix(0, ncol(mat), ncol(mat))
  colnames(MCnemarC) <- colnames(mat)
  rownames(MCnemarC) <- colnames(mat)
  
  for (i in 1:ncol(mat)) {
    for (j in 1:ncol(mat)) {
      if (i != j) {
        tab <- matrix(c(Xij[i, j], Xi[i]-Xij[i, j],Xi[j]-Xij[i, j],  301-Xi[i]-Xi[j]+Xij[i, j]), 
                      nrow = 2, byrow = TRUE)
        MCnemarC[i, j] <- round(p.adjust(mcnemar.test(tab)$statistic, method = "holm"),2)
      }
    }
  }
  
  return(list(count = Xij, sri = SRI, mcnP = MCnemarP, mcnChi = MCnemarC))
}

#Convert to long format for plotting
plot_matrix <- function(Xij, SRI, genus_name) {
  
  # Reorder hosts (same order for both axes)
  ord <- seriate(dist(Xij), method = "OLO")
  order_idx <- get_order(ord)
  
  Xij <- Xij[order_idx, order_idx]
  SRI <- SRI[order_idx, order_idx]

  # Convert to long format AFTER reordering
  df_counts <- as.data.frame(as.table(as.matrix(Xij)))
  colnames(df_counts) <- c("Host1", "Host2", "Count")

  df_sri <- as.data.frame(as.table(as.matrix(SRI)))
  colnames(df_sri) <- c("Host1", "Host2", "SRI")


  df <- merge(df_counts, df_sri, by = c("Host1", "Host2"))

  # Ensure identical factor ordering for x and y
  host_levels <- colnames(Xij)
  
  df$Host1 <- factor(df$Host1, levels = host_levels)
  df$Host2 <- factor(df$Host2, levels = host_levels)

  # Triangle classification
  df <- df %>%
    mutate(
      i = as.numeric(Host1),
      j = as.numeric(Host2),
      lower = i > j,
      upper = i < j,
      diag  = i == j
    )

  ggplot() +

    # LOWER TRIANGLE → COUNTS
    geom_tile(
      data = subset(df, lower),
      aes(Host1, Host2, fill = Count),
      color = "white"
    ) +
    geom_text(
      data = subset(df, lower),
      aes(Host1, Host2, label = Count),
      size = 3
    ) +

    scale_fill_gradient(
      low = "white",
      high = "#C11007",
      name = "Count"
    ) +

    # New scale for SRI
    ggnewscale::new_scale_fill() +

    # UPPER TRIANGLE → SRI
    geom_tile(
      data = subset(df, upper),
      aes(Host1, Host2, fill = SRI),
      color = "white"
    ) +
    geom_text(
      data = subset(df, upper),
      aes(x = Host1, y = Host2, label = round(SRI, 2)),
      size = 3
    ) +

    scale_fill_gradient(
      low = "white",
      high = "#075BC1",
      name = "SRI"
    ) +

    # DIAGONAL → COUNTS (highlighted)
    geom_tile(
      data = subset(df, diag),
      aes(Host1, Host2),
      fill = "grey85",
      color = "black"
    ) +
    geom_text(
      data = subset(df, diag),
      aes(x = Host1, y = Host2, label = Count),
      size = 3,
      fontface = "bold"
    ) +
    
    scale_x_discrete(limits = host_levels) +
    
    scale_y_discrete(limits = host_levels) +

    theme_minimal() +
    labs(
      title = paste0(
        "Host group feeding overlap ", genus_name,
        " (N = ", MQ_counts[genus_name], ")"
      ),
      x = "Host group",
      y = "Host group"
    ) +
    theme(
      axis.text.x = element_text(angle = 45, hjust = 1),
      panel.grid = element_blank()
    )
}

# Create empty list to store plots
matrix_plots <- list()

# Create visual matrix for each genus
for (genus in names(genus_list)) {

  mats <- build_host_matrices(genus_list[[genus]])

  p <- plot_matrix(mats$count, mats$sri, genus)

  matrix_plots[[genus]] <- p

  print(p)
  
  print(paste0("MCnemar Chi-square for ", genus))
  print(mats$mcnChi)
  print(paste0("MCnemar holm adjusted p-value for ", genus))
  print(mats$mcnP)
  
  ggsave(
    filename = paste0("HostFeedingMatrix_", genus, ".svg"),
    plot = p,
    width = 10,
    height = 6)
}
```

```
## [1] "MCnemar Chi-square for Aedes"
##       Human Shrew
## Human  0.00  2.25
## Shrew  2.25  0.00
## [1] "MCnemar holm adjusted p-value for Aedes"
##       Human Shrew
## Human 1.000 0.134
## Shrew 0.134 1.000
```

```
## [1] "MCnemar Chi-square for Anopheles"
##                 Domestic mammal
## Domestic mammal               0
## [1] "MCnemar holm adjusted p-value for Anopheles"
##                 Domestic mammal
## Domestic mammal               1
```

```
## [1] "MCnemar Chi-square for Culex"
##                       Human  Crow Water bird Commensal rodent Wild mammal
## Human                  0.00 94.88     164.39           182.40      199.00
## Crow                  94.88  0.00      20.34            26.27       45.45
## Water bird           164.39 20.34       0.00             0.14        6.05
## Commensal rodent     182.40 26.27       0.14             0.00        4.27
## Wild mammal          199.00 45.45       6.05             4.27        0.00
## Non commensal rodent  96.75  1.06      32.47            40.13       55.71
## Domestic bird        187.13 28.92       0.32             0.00        3.06
## Other bird           185.80 37.77       3.68             1.89        0.10
## Shrew                173.85 17.75       0.03             0.52        7.68
## Domestic mammal      180.08 29.78       1.14             0.19        1.79
## Non human primate    202.00 50.16      11.53             8.64        0.80
##                      Non commensal rodent Domestic bird Other bird  Shrew
## Human                               96.75        187.13     185.80 173.85
## Crow                                 1.06         28.92      37.77  17.75
## Water bird                          32.47          0.32       3.68   0.03
## Commensal rodent                    40.13          0.00       1.89   0.52
## Wild mammal                         55.71          3.06       0.10   7.68
## Non commensal rodent                 0.00         36.91      49.32  33.39
## Domestic bird                       36.91          0.00       1.39   0.89
## Other bird                          49.32          1.39       0.00   5.04
## Shrew                               33.39          0.89       5.04   0.00
## Domestic mammal                     41.81          0.05       0.56   1.75
## Non human primate                   62.13          7.69       2.29  13.47
##                      Domestic mammal Non human primate
## Human                         180.08            202.00
## Crow                           29.78             50.16
## Water bird                      1.14             11.53
## Commensal rodent                0.19              8.64
## Wild mammal                     1.79              0.80
## Non commensal rodent           41.81             62.13
## Domestic bird                   0.05              7.69
## Other bird                      0.56              2.29
## Shrew                           1.75             13.47
## Domestic mammal                 0.00              5.82
## Non human primate               5.82              0.00
## [1] "MCnemar holm adjusted p-value for Culex"
##                      Human  Crow Water bird Commensal rodent Wild mammal
## Human                    1 0.000      0.000            0.000       0.000
## Crow                     0 1.000      0.000            0.000       0.000
## Water bird               0 0.000      1.000            0.710       0.014
## Commensal rodent         0 0.000      0.710            1.000       0.039
## Wild mammal              0 0.000      0.014            0.039       1.000
## Non commensal rodent     0 0.303      0.000            0.000       0.000
## Domestic bird            0 0.000      0.571            1.000       0.080
## Other bird               0 0.000      0.055            0.169       0.752
## Shrew                    0 0.000      0.864            0.472       0.006
## Domestic mammal          0 0.000      0.286            0.663       0.181
## Non human primate        0 0.000      0.001            0.003       0.371
##                      Non commensal rodent Domestic bird Other bird Shrew
## Human                               0.000         0.000      0.000 0.000
## Crow                                0.303         0.000      0.000 0.000
## Water bird                          0.000         0.571      0.055 0.864
## Commensal rodent                    0.000         1.000      0.169 0.472
## Wild mammal                         0.000         0.080      0.752 0.006
## Non commensal rodent                1.000         0.000      0.000 0.000
## Domestic bird                       0.000         1.000      0.239 0.345
## Other bird                          0.000         0.239      1.000 0.025
## Shrew                               0.000         0.345      0.025 1.000
## Domestic mammal                     0.000         0.831      0.453 0.186
## Non human primate                   0.000         0.006      0.131 0.000
##                      Domestic mammal Non human primate
## Human                          0.000             0.000
## Crow                           0.000             0.000
## Water bird                     0.286             0.001
## Commensal rodent               0.663             0.003
## Wild mammal                    0.181             0.371
## Non commensal rodent           0.000             0.000
## Domestic bird                  0.831             0.006
## Other bird                     0.453             0.131
## Shrew                          0.186             0.000
## Domestic mammal                1.000             0.016
## Non human primate              0.016             1.000
```

```
## [1] "MCnemar Chi-square for Mansonia"
##                      Human Non commensal rodent Commensal rodent Water bird
## Human                    0                    0                0          0
## Non commensal rodent     0                    0                0          0
## Commensal rodent         0                    0                0        NaN
## Water bird               0                    0              NaN          0
## [1] "MCnemar holm adjusted p-value for Mansonia"
##                      Human Non commensal rodent Commensal rodent Water bird
## Human                    1                    1                1          1
## Non commensal rodent     1                    1                1          1
## Commensal rodent         1                    1                1        NaN
## Water bird               1                    1              NaN          1
```

```
## The matrix visualizes host group feeding overlap: diagonal values indicate how many mosquitoes fed on each host group, the lower triangle shows the number of mosquitoes that fed on both host groups, and the upper triangle shows the strength of association between host groups (SRI).
```

#### 5.2 Culex network analysis

##### 5.2.1 Sitesplitting

```
# Split data by mosquito genus Culex and site
Site_list <- split(genus_list$Culex, genus_list$Culex$Site)

# Count the number of Culex for each site
Site_counts <- c(
  Rural     = length(unique(Site_list$Rural$MQ)),
  Suburban = length(unique(Site_list$Suburban$MQ)),
  Urban     = length(unique(Site_list$Urban$MQ))
)
```

###### 5.2.1.1 Host–host network construction

```
# Create networks
host_Sitenetworks <- lapply(Site_list, build_host_network_SRI)

# Store genus names
site_names <- names(host_Sitenetworks)

# Create network plots for each site
for (site in site_names) {

  res <- host_Sitenetworks[[site]]
  g <- res$graph
  mat <- res$matrix   # <-- matrix available
  
  props <- host_mosquito_prop(Site_list[[site]])

  V(g)$prop <- props$prop[match(V(g)$name, props$HostGroup)]
  V(g)$prop[is.na(V(g)$prop)] <- 0

  # Edge weights already SRI
  E(g)$weight <- as.numeric(E(g)$weight)
  
  # Plot
  p <- ggraph(g, layout = "fr") +
    geom_edge_link(
      aes(edge_color = weight),
      alpha = 0.8,
      width = 2.5
    ) +
    geom_node_point(
      aes(size = prop),
      colour = "#C11007"
    ) +
    geom_node_text(
      aes(label = name),
      repel = TRUE
    ) +
    scale_edge_color_gradient(low = "white",
      high = "#075BC1",limits = c(0, max(E(g)$weight))) +
                              
    labs(
      title = paste0("Culex feeding network in ", site,
                     " site (N= ", Site_counts[site], ")"),
      edge_color = "SRI",
      size = "Host group proportion"
    ) +
    theme_void()

  network_plots[[site]] <- p
  print(p)
  
  ggsave(
    filename = paste0("FeedingNetwork_Culex_", site, ".svg"),
    plot = p,
    width = 10,
    height = 6)
}
```

###### 5.2.1.2 Host-host matrix construction

```
#Convert to long format for plotting
SitePlot_matrix <- function(Xij, SRI, site_name) {
  
  # Reorder hosts (same order for both axes)
  ord <- seriate(dist(Xij), method = "OLO")
  order_idx <- get_order(ord)
  
  Xij <- Xij[order_idx, order_idx]
  SRI <- SRI[order_idx, order_idx]

  # Convert to long format AFTER reordering
  df_counts <- as.data.frame(as.table(as.matrix(Xij)))
  colnames(df_counts) <- c("Host1", "Host2", "Count")

  df_sri <- as.data.frame(as.table(as.matrix(SRI)))
  colnames(df_sri) <- c("Host1", "Host2", "SRI")


  df <- merge(df_counts, df_sri, by = c("Host1", "Host2"))

  # Ensure identical factor ordering for x and y
  host_levels <- colnames(Xij)
  
  df$Host1 <- factor(df$Host1, levels = host_levels)
  df$Host2 <- factor(df$Host2, levels = host_levels)

  # Triangle classification
  df <- df %>%
    mutate(
      i = as.numeric(Host1),
      j = as.numeric(Host2),
      lower = i > j,
      upper = i < j,
      diag  = i == j
    )

  ggplot() +

    # LOWER TRIANGLE → COUNTS
    geom_tile(
      data = subset(df, lower),
      aes(Host1, Host2, fill = Count),
      color = "white"
    ) +
    geom_text(
      data = subset(df, lower),
      aes(Host1, Host2, label = Count),
      size = 3
    ) +

    scale_fill_gradient(
      low = "white",
      high = "#C11007",
      name = "Count"
    ) +

    # New scale for SRI
    ggnewscale::new_scale_fill() +

    # UPPER TRIANGLE → SRI
    geom_tile(
      data = subset(df, upper),
      aes(Host1, Host2, fill = SRI),
      color = "white"
    ) +
    geom_text(
      data = subset(df, upper),
      aes(x = Host1, y = Host2, label = round(SRI, 2)),
      size = 3
    ) +

    scale_fill_gradient(
      low = "white",
      high = "#075BC1",
      name = "SRI"
    ) +

    # DIAGONAL → COUNTS (highlighted)
    geom_tile(
      data = subset(df, diag),
      aes(Host1, Host2),
      fill = "grey85",
      color = "black"
    ) +
    geom_text(
      data = subset(df, diag),
      aes(x = Host1, y = Host2, label = Count),
      size = 3,
      fontface = "bold"
    ) +
    
    scale_x_discrete(limits = host_levels) +
    
    scale_y_discrete(limits = host_levels) +

    theme_minimal() +
    labs(
      title = paste0(
        "Culex host group feeding overlap in ", site_name,
        " site (N = ", Site_counts[site_name], ")"
      ),
      x = "Host group",
      y = "Host group"
    ) +
    theme(
      axis.text.x = element_text(angle = 45, hjust = 1),
      panel.grid = element_blank()
    )
}

# Create visual matrix for each genus
for (site in names(Site_list)) {

  mats <- build_host_matrices(Site_list[[site]])

  p <- SitePlot_matrix(mats$count, mats$sri, site)

  matrix_plots[[site]] <- p

  print(p)
  
  ggsave(
    filename = paste0("HostFeedingMatrix_Culex_", site, ".svg"),
    plot = p,
    width = 10,
    height = 6)
}
```

##### 5.2.2 Season splitting

```
# Split data by mosquito genus Culex and season
Season_list <- split(genus_list$Culex, genus_list$Culex$Season)

# Count the number of Culex for each year
Season_counts <- c(
  Wet     = length(unique(Season_list$Wet$MQ)),
  Dry = length(unique(Season_list$Dry$MQ))
)
```

###### 5.2.2.1 Host–host network construction

```
# Create networks
host_Seasonnetworks <- lapply(Season_list, build_host_network_SRI)

# Store seson names
season_names <- names(host_Seasonnetworks)

# Create network plots for each season
for (season in season_names) {

  res <- host_Seasonnetworks[[season]]
  g <- res$graph
  mat <- res$matrix   # <-- matrix available
  
  props <- host_mosquito_prop(Season_list[[season]])

  V(g)$prop <- props$prop[match(V(g)$name, props$HostGroup)]
  V(g)$prop[is.na(V(g)$prop)] <- 0

  # Edge weights already SRI
  E(g)$weight <- as.numeric(E(g)$weight)

  # Plot
  p <- ggraph(g, layout = "fr") +
    geom_edge_link(
      aes(edge_color = weight),
      alpha = 0.8,
      width = 2.5
    ) +
    geom_node_point(
      aes(size = prop),
      colour = "#C11007"
    ) +
    geom_node_text(
      aes(label = name),
      repel = TRUE
    ) +
    scale_edge_color_gradient(low = "white",
      high = "#075BC1",limits = c(0, max(E(g)$weight))) +
                              
    labs(
      title = paste0("Culex feeding network in ", season,
                     " season (N= ", Season_counts[season], ")"),
      edge_color = "SRI",
      size = "Host group proportion"
    ) +
    theme_void()

  network_plots[[season]] <- p
  print(p)
  
  ggsave(
    filename = paste0("FeedingNetwork_Culex_", season, ".svg"),
    plot = p,
    width = 10,
    height = 6)
}
```

###### 5.2.2.2 Host-host matrix construction

```
#Convert to long format for plotting
SeasonPlot_matrix <- function(Xij, SRI, season_name) {
  
  # Reorder hosts (same order for both axes)
  ord <- seriate(dist(Xij), method = "OLO")
  order_idx <- get_order(ord)
  
  Xij <- Xij[order_idx, order_idx]
  SRI <- SRI[order_idx, order_idx]

  # Convert to long format AFTER reordering
  df_counts <- as.data.frame(as.table(as.matrix(Xij)))
  colnames(df_counts) <- c("Host1", "Host2", "Count")

  df_sri <- as.data.frame(as.table(as.matrix(SRI)))
  colnames(df_sri) <- c("Host1", "Host2", "SRI")


  df <- merge(df_counts, df_sri, by = c("Host1", "Host2"))

  # Ensure identical factor ordering for x and y
  host_levels <- colnames(Xij)
  
  df$Host1 <- factor(df$Host1, levels = host_levels)
  df$Host2 <- factor(df$Host2, levels = host_levels)

  # Triangle classification
  df <- df %>%
    mutate(
      i = as.numeric(Host1),
      j = as.numeric(Host2),
      lower = i > j,
      upper = i < j,
      diag  = i == j
    )

  ggplot() +

    # LOWER TRIANGLE → COUNTS
    geom_tile(
      data = subset(df, lower),
      aes(Host1, Host2, fill = Count),
      color = "white"
    ) +
    geom_text(
      data = subset(df, lower),
      aes(Host1, Host2, label = Count),
      size = 3
    ) +

    scale_fill_gradient(
      low = "white",
      high = "#C11007",
      name = "Count"
    ) +

    # New scale for SRI
    ggnewscale::new_scale_fill() +

    # UPPER TRIANGLE → SRI
    geom_tile(
      data = subset(df, upper),
      aes(Host1, Host2, fill = SRI),
      color = "white"
    ) +
    geom_text(
      data = subset(df, upper),
      aes(x = Host1, y = Host2, label = round(SRI, 2)),
      size = 3
    ) +

    scale_fill_gradient(
      low = "white",
      high = "#075BC1",
      name = "SRI"
    ) +

    # DIAGONAL → COUNTS (highlighted)
    geom_tile(
      data = subset(df, diag),
      aes(Host1, Host2),
      fill = "grey85",
      color = "black"
    ) +
    geom_text(
      data = subset(df, diag),
      aes(x = Host1, y = Host2, label = Count),
      size = 3,
      fontface = "bold"
    ) +
    
    scale_x_discrete(limits = host_levels) +
    
    scale_y_discrete(limits = host_levels) +

    theme_minimal() +
    labs(
      title = paste0(
        "Culex host group feeding overlap in ", season_name,
        " season (N = ", Season_counts[season_name], ")"
      ),
      x = "Host group",
      y = "Host group"
    ) +
    theme(
      axis.text.x = element_text(angle = 45, hjust = 1),
      panel.grid = element_blank()
    )
}

# Create visual matrix for each genus
for (season in names(Season_list)) {

  mats <- build_host_matrices(Season_list[[season]])

  p <- SeasonPlot_matrix(mats$count, mats$sri, season)

  matrix_plots[[season]] <- p

  print(p)
  
  ggsave(
    filename = paste0("HostFeedingMatrix_Culex_", season, ".svg"),
    plot = p,
    width = 10,
    height = 6)
}
```

### 6 Human feedings

#### 6.1 Filtering to human feedings

```
HumanCulex_data <- FilterHost %>%
  filter(VectorGenus == "Culex") %>%                # keep only Culex
  group_by(MQ) %>%
  filter(any(HostGroup == "Human" & HostProportion > 0)) %>%  # keep MQ with human feeding
  ungroup()

# Define full host set ONCE
all_hosts <- unique(HumanCulex_data$HostGroup)

presence_df <- HumanCulex_data %>%
  
  mutate(Presence = ifelse(HostProportion > 0, 1, 0)) %>%
  
  select(MQ, Site, Season, Host = HostGroup, Presence) %>%
  
  group_by(MQ, Site, Season, Host) %>%
  summarise(Presence = max(Presence), .groups = "drop") %>%
  
  # group first (preserve nesting)
  group_by(MQ, Site, Season) %>%
  
  # complete using FULL host list
  complete(
    Host = all_hosts,
    fill = list(Presence = 0)
  ) %>%
  
  ungroup() %>%
  filter(Host != "Human")

# Number of mosquitoes having fed on human
HumanMQ <- length(unique(presence_df$MQ))

# Table of counts for each host season & site, used Crows to filter since for every host mosquito combination exists
HumanMQsubdivision <- table(presence_df$Season[presence_df$Host== "Crow"], presence_df$Site[presence_df$Host== "Crow"])
```

```
## Total number of mosquitoes that have fed on humans
```

```
## [1] 205
```

```
## Total number of mosquitoes having fed on humans according to season and site
```

#### 6.2 Overall host model

```
# Make an overall host model
model_host <- glmer(
  Presence ~ Host + (1 | MQ),
  data = presence_df,
  family = binomial
)

drop1(model_host, test = "Chisq")
```

|  | npar | AIC | LRT | Pr(Chi) |
| --- | --- | --- | --- | --- |
|  | NA | 674.1455 | NA | NA |
| Host | 8 | 746.3008 | 88.15528 | 0 |

```
summary(model_host)
```

```
## Generalized linear mixed model fit by maximum likelihood (Laplace
##   Approximation) [glmerMod]
##  Family: binomial  ( logit )
## Formula: Presence ~ Host + (1 | MQ)
##    Data: presence_df
## 
##       AIC       BIC    logLik -2*log(L)  df.resid 
##     674.1     729.3    -327.1     654.1      1835 
## 
## Scaled residuals: 
##     Min      1Q  Median      3Q     Max 
## -0.6125 -0.2185 -0.1794 -0.1015 14.1432 
## 
## Random effects:
##  Groups Name        Variance Std.Dev.
##  MQ     (Intercept) 0.4573   0.6762  
## Number of obs: 1845, groups:  MQ, 205
## 
## Fixed effects:
##                          Estimate Std. Error z value Pr(>|z|)    
## (Intercept)               -3.2816     0.3758  -8.732  < 2e-16 ***
## HostCrow                   0.3950     0.4496   0.879   0.3797    
## HostDomestic bird          0.1108     0.4745   0.234   0.8154    
## HostDomestic mammal       -1.1377     0.6764  -1.682   0.0926 .  
## HostNon commensal rodent   1.6917     0.3919   4.316 1.59e-05 ***
## HostNon human primate     -2.2533     1.0608  -2.124   0.0337 *  
## HostShrew                  0.3087     0.4566   0.676   0.4990    
## HostWater bird            -1.1449     0.6781  -1.688   0.0913 .  
## HostWild mammal           -0.8430     0.6118  -1.378   0.1682    
## ---
## Signif. codes:  0 '***' 0.001 '**' 0.01 '*' 0.05 '.' 0.1 ' ' 1
## 
## Correlation of Fixed Effects:
##             (Intr) HstCrw HstDmb HstDmm HstNcr HstNhp HstShr HstWtb
## HostCrow    -0.703                                                 
## HstDmstcbrd -0.663  0.553                                          
## HstDmstcmmm -0.460  0.388  0.368                                   
## HstNncmmnsr -0.843  0.671  0.635  0.444                            
## HstNnhmnprm -0.292  0.247  0.234  0.165  0.283                     
## HostShrew   -0.691  0.575  0.545  0.382  0.661  0.244              
## HostWatrbrd -0.459  0.387  0.367  0.258  0.443  0.164  0.381       
## HostWldmmml -0.509  0.429  0.407  0.286  0.491  0.182  0.423  0.285
## optimizer (Nelder_Mead) convergence code: 0 (OK)
## Model failed to converge with max|grad| = 0.00978226 (tol = 0.002, component 1)
##   See ?lme4::convergence and ?lme4::troubleshooting.
```

```
# Emmeans of the host model
emm <- emmeans(model_host, ~ Host)

# Make compact letter display showing significant differences
QQ <- as.data.frame(cld(emm , type = 'response', Letters = letters, sort = FALSE))
QQ$.group <- gsub(" ", "", QQ$.group)
QQ <- QQ %>%
  mutate(across(c(prob, SE, asymp.LCL, asymp.UCL),
                ~ round(.x * 100, 1)))

#Alter grouping since only 1 host is significantly different
QQ$.group <- ifelse(QQ$.group == "a", "", "*")

# Create plot
CoFeedHost <- ggplot(data = QQ ,aes(x = Host, y = prob)) +
  geom_point(position = position_dodge(width = 0.7)) +
  geom_errorbar(aes(ymin = asymp.LCL, ymax = asymp.UCL),
                width = 0.2, 
                position = position_dodge(width = 0.7))+
  geom_text(aes(y = asymp.UCL + 1, label = .group),
            position = position_dodge(width = 0.7), 
            size = 4, 
            show.legend = FALSE) +
  labs(x = "Host group", y = "Co-feeding probability with 95% CI (%)",
      title = expression("Human bloodfed " * italic("Culex")*"(N=205)"))+
  theme_bw()+
  theme(
    axis.text.x = element_text(angle = 45, hjust = 1),
    axis.title = element_text(size = 14),       # x and y axis titles
    axis.text  = element_text(size = 12),       # x and y axis tick labels
    legend.title = element_text(size = 14),     # legend title
    legend.text  = element_text(size = 12)      # legend items
  )

ggsave(
    filename = "Co-feedingHuman_Host.svg",
    plot = CoFeedHost,
    width = 10,
    height = 6)
```

```
## Estimated Marginal means of the host model
```

```
## Odds ratio of the Estimated Marginal means of the host model with p values
```

#### 6.3 Full interaction model

```
# All interactions and variables
full_model<- glmer(Presence ~ Host + Season + Site + Host:Season + Host:Site + Season:Site + (1 | MQ),
  data = presence_df,
  family = binomial
)
drop1(full_model, test = "Chisq")
```

|  | npar | AIC | LRT | Pr(Chi) |
| --- | --- | --- | --- | --- |
|  | NA | 659.7152 | NA | NA |
| Host:Season | 8 | 677.5899 | 33.874715 | 0.0000428 |
| Host:Site | 16 | 652.1994 | 24.484187 | 0.0794498 |
| Season:Site | 2 | 658.3215 | 2.606362 | 0.2716662 |

```
# Only significant ones
full_model2<- glmer(Presence ~ Host + Season + Site + Host:Season + (1 | MQ),
  data = presence_df,
  family = binomial
)
drop1(full_model2, test = "Chisq")
```

|  | npar | AIC | LRT | Pr(Chi) |
| --- | --- | --- | --- | --- |
|  | NA | 648.8841 | NA | NA |
| Site | 2 | 652.4514 | 7.567283 | 0.0227397 |
| Host:Season | 8 | 664.0890 | 31.204953 | 0.0001292 |

#### 6.4 Host site model

```
# Make host site model even though it has been shown to not be significant
model_site <- glmer(Presence ~ Host * Site + (1 | MQ), data = presence_df, family = binomial)

drop1(model_site, test = "Chisq")
```

|  | npar | AIC | LRT | Pr(Chi) |
| --- | --- | --- | --- | --- |
|  | NA | 682.4459 | NA | NA |
| Host:Site | 16 | 672.5036 | 22.0577 | 0.1413381 |

```
# Emmeans of the host-site model
emm <- emmeans(model_site, ~ Host | Site)

# Make compact letter display showing significant differences
QQ <- as.data.frame(cld(emm , type = 'response', Letters = letters, sort = FALSE))
QQ$.group <- gsub(" ", "", QQ$.group)
QQ <- QQ %>%
  mutate(across(c(prob, SE, asymp.LCL, asymp.UCL),~ round(.x * 100, 1)))%>%
  mutate(asymp.UCL = ifelse(asymp.UCL >= 100, 0, asymp.UCL))

# Create plot
CoFeedHostSite <- ggplot(data = QQ, aes(x = Host, y = prob, color = Site)) +
  geom_point(position = position_dodge(width = 0.7)) +
  geom_errorbar(aes(ymin = asymp.LCL, ymax = asymp.UCL),
                width = 0.2, 
                position = position_dodge(width = 0.7)) +
  geom_text(aes(y = asymp.UCL + 4, label = .group),position = position_dodge(width = 0.7), size = 4, show.legend = FALSE) +
    facet_wrap(~Site, ncol =1)+
  labs(
    x = "Host group",
    y = "Co-feeding probability with 95% CI (%)",
    title = expression("Human bloodfed " * italic("Culex") * "(N=205)")
  ) +
  theme_bw() +
  theme(
    axis.text.x = element_text(angle = 45, hjust = 1),
    axis.title = element_text(size = 14),
    axis.text  = element_text(size = 12),
    legend.position = "top" 
  )
```

```
## Estimated Marginal means of the host-site model
```

```
## Odds ratio of the Estimated Marginal means of the host-site model with p values
```

#### 6.5 Host Season model

```
# Make model
model_season <- glmer(
  Presence ~ Host * Season + (1 | MQ),
  data = presence_df,
  family = binomial)

drop1(model_season, test = "Chisq")
```

|  | npar | AIC | LRT | Pr(Chi) |
| --- | --- | --- | --- | --- |
|  | NA | 652.4514 | NA | NA |
| Host:Season | 8 | 667.6331 | 31.18175 | 0.0001304 |

```
# Emmeans of the host-season model
emm <- emmeans(model_season, ~ Host * Season)

# Determine the significant differences but only for either shared host or shared season
pairs_filtered <- as.data.frame(pairs(emm, type = "response")) %>%
  separate(contrast, into = c("grp1", "grp2"), sep = " / ") %>%
  separate(grp1, into = c("Host1", "Season1"), sep = " ") %>%
  separate(grp2, into = c("Host2", "Season2"), sep = " ") %>%
  filter(
    Host1 == Host2 | Season1 == Season2
  ) %>%
  filter( p.value <=0.05)

# Make compact letter display showing significant differences
QQ <- as.data.frame(cld(emm , type = 'response', Letters = letters, sort = FALSE))
QQ$.group <- gsub(" ", "", QQ$.group)
QQ <- QQ %>%
  mutate(across(c(prob, SE, asymp.LCL, asymp.UCL),~ round(.x * 100, 1)))%>%
  mutate(asymp.UCL = ifelse(asymp.UCL >= 100, 0, asymp.UCL))

# Create plot
CoFeedHostSeason <- ggplot(data = QQ ,aes(x = Host, y = prob, color = Season)) +
  geom_point(position = position_dodge(width = 0.7)) +
  geom_errorbar(aes(ymin = asymp.LCL, ymax = asymp.UCL),
                width = 0.2, 
                position = position_dodge(width = 0.7))+
  #geom_text(aes(y = asymp.UCL + 2, label = .group), position = position_dodge(width = 0.7), size = 4, show.legend = FALSE) +
  labs(x = "Host group", y = "Co-feeding probability with 95% CI (%)",
      title = expression("Human bloodfed " * italic("Culex")*"(N=205)"))+
  theme_bw()+
  theme(
    axis.text.x = element_text(angle = 45, hjust = 1),
    axis.title = element_text(size = 14),       # x and y axis titles
    axis.text  = element_text(size = 12),       # x and y axis tick labels
    legend.title = element_text(size = 14),     # legend title
    legend.text  = element_text(size = 12)      # legend items
  )

sig_labels <- data.frame(
  x = c(4.55, 7.15),
  y = c(54, 58),
  label = c("p=0.038", "p=0.045")
)

CoFeedHostSeason <- CoFeedHostSeason +
  geom_segment(aes(x = 4.15, xend = 5.1, 
                   y = 52, yend = 52), color = "black") +
  geom_segment(aes(x = 5.2, xend = 9.15, 
                   y = 56, yend = 56), color = "black") +

  # vertical ticks (optional but nicer)
  geom_segment(aes(x = 4.15, xend = 4.15, 
                   y = 10, yend = 52), color = "black", linetype = 2) +
  geom_segment(aes(x = 5.1, xend = 5.1, 
                   y = 25, yend = 52), color = "black", linetype = 2) +

  geom_segment(aes(x = 5.2, xend = 5.2, 
                   y = 25, yend = 56), color = "black", linetype = 2) +
  geom_segment(aes(x = 9.15, xend = 9.15, 
                   y = 10, yend = 56), color = "black", linetype = 2) +
  geom_text(data = sig_labels,
            aes(x = x, y = y, label = label),
            inherit.aes = FALSE,
            color = "black")

ggsave(
    filename = "Co-feedingHuman_HostSeason.svg",
    plot = CoFeedHostSeason,
    width = 10,
    height = 6)
```

```
## Estimated Marginal means of the host-season model
```

```
## Odds ratio of the Estimated Marginal means of the host-season model with p values
```

```
## Odds ratio of the Estimated Marginal means of the host-season model with p values for only the relevant significant interactions
```

| Host1 | Season1 | Host2 | Season2 | odds.ratio | SE | df | null | z.ratio | p.value |
| --- | --- | --- | --- | --- | --- | --- | --- | --- | --- |
| Non | commensal | Non | human | 100.2874 | 114.7667 | Inf | 1 | 4.026673 | 0.007158 |
